# *UBA1-CDK16*: A Sex-Specific Chimeric RNA and Its Role in Immune Sexual Dimorphism

**DOI:** 10.1101/2024.02.13.580120

**Authors:** Xinrui Shi, Loryn Facemire, Sandeep Singh, Shailesh Kumar, Robert Cornelison, Chen Liang, Fujun Qin, Aiqun Liu, Shitong Lin, Yue Tang, Justin Elfman, Thomas Manley, Timothy Bullock, Doris M. Haverstick, Peng Wu, Hui Li

**Author notes:** Corresponding author., Web: http://lilab.medicine.virginia.edu.

## Abstract

RNA processing mechanisms, such as alternative splicing and RNA editing, have been recognized as critical means to expand the transcriptome. Chimeric RNAs formed by intergenic splicing provide another potential layer of RNA diversification. By analyzing a large set of RNA-Seq data and validating results in over 1,200 blood samples, we identified *UBA1-CDK16*, a female-specific chimeric transcript. Intriguingly, both parental genes, are expressed in males and females. Mechanistically, *UBA1-CDK16* is produced by cis-splicing between the two adjacent X-linked genes, originating from the inactive X chromosome. A female-specific chromatin loop, formed between the junction sites, facilitates the alternative splicing of its readthrough precursor. This unique chimeric transcript exhibits evolutionary conservation, evolving to be female-specific from non-human primates to humans. Furthermore, our investigation reveals that *UBA1-CDK16* is enriched in the myeloid lineage and plays a regulatory role in myeloid differentiation. Notably, female COVID-19 patients who tested negative for this chimeric transcript displayed higher counts of neutrophils, highlighting its potential role in disease pathogenesis. These findings support the notion that chimeric RNAs represent a new repertoire of transcripts that can be regulated independently from the parental genes, and a new class of RNA variance with potential implications in sexual dimorphism and immune responses.

## Introduction

The complexity of an organism is not completely represented by the genome size and the number of genes^1^. Alternative splicing^2^, along with other processes such as RNA editing^3^, are known to expand the functional genome. Chimeric RNAs are fusion transcripts composed of sequences transcribed from disparate parental genes^4,5^. Traditionally, they were thought to be unique features of cancer cells^6–9^. However, an increasing number of chimeric RNAs have been reported in non-cancerous cells and tissues^10–12^, suggesting their role in diversifying the transcriptome^4,13^.

Chimeric RNAs formation encompasses diverse mechanisms. Traditionally, these RNAs are considered the results of fusion genes arising from chromosomal rearrangements such as translocations, deletions, and inversions, often linked to cancer pathogenesis^14,15^. However, at least two other mechanisms are now known to generate chimeric RNAs, independent of such chromosome rearrangements. These mechanisms include trans-splicing^10^, involving the fusion of exons from different transcripts, and cis-splicing between adjacent genes (cis-SAGe)^16,17^, which occurs when the transcriptional machinery continues beyond the typical gene termination site and enters the adjacent gene. However, the precise molecular mechanisms driving trans-splicing and cis-SAGe remain largely unknown.

While men and women share identical genetic information across the majority of their genomes, they exhibit distinct sex-specific characteristics such as physical performance, behavior, and physiological traits. The sexual dimorphism is determined by various factors, including sex chromosomes, sex hormones, sexual transcriptomic variations, and sexual epigenomic dynamics^18,19^. In humans, sexual dimorphism is notably pronounced in immune responses, leading to differential responses to diseases^20^. Studies have shown that females often mount stronger immune responses to infections than males and tend to have higher prevalence of autoimmune diseases^21,22^.

To delve deeper into the landscape of the sexome, we first explored the expression and regulation of chimeric RNAs in male and female tissues using the Genotype-Tissue Expression (GTEx) Project dataset^23,24^. Using this dataset, we identified a novel X-linked chimeric RNA, *UBA1-CDK16,* almost exclusively expressed in female blood samples. Subsequently, we rigorously validated this discovery in an extensive collection of clinical samples. Mechanistically, we found that the chimeric RNA arises as a product of cis-SAGe from the inactive X chromosome. We elucidate the existence of a sex-specific chromatin loop formed between the junction splicing sites, which significantly contributes to cis-SAGe. Furthermore, we unraveled the functional role of this chimeric RNA as a pivotal checkpoint inhibitor during myeloid differentiation. This study presents a novel perspective on sexual dimorphism, underscoring the crucial role of a sex-specific chimeric RNA emerging from Xi-specific chromatin structures. Through this discovery, we illuminate the involvement of chimeric RNAs in the context of sex-biased immunity and provide profound implications for sex-biased diseases.

## Results

### Identification of a female-specific *UBA1-CDK16* chimeric RNA in human blood

Using EricScript^25^, we examined the chimeric RNA transcriptome within the Genotype-Tissue Expression (GTEx) database. GTEx includes RNA-seq data from 44 non-diseased tissue sites across healthy individuals^23,24^. In prioritizing sample accessibility for downstream validation, we primarily focused on whole blood tissue type. A total of 57,063 chimeric RNAs were identified from 426 whole blood RNA-seq dataset in GTEx, from 157 women and 269 men. Chimeric RNAs were classified based on the junction site location: if the junction site fell within 2bp of the canonical exon boundary, the junction was annotated as “E” (edge of exon); otherwise, it was annotated as “M” (middle of exon). To reduce potential false positives identified by EricScript, we filtered out “MM” fusions^26^, and focused on recurrent chimeras found in more than five individuals, which resulted in the final selection of 1,158 chimeric RNAs (**Fig. 1a**).

**Figure 1.**
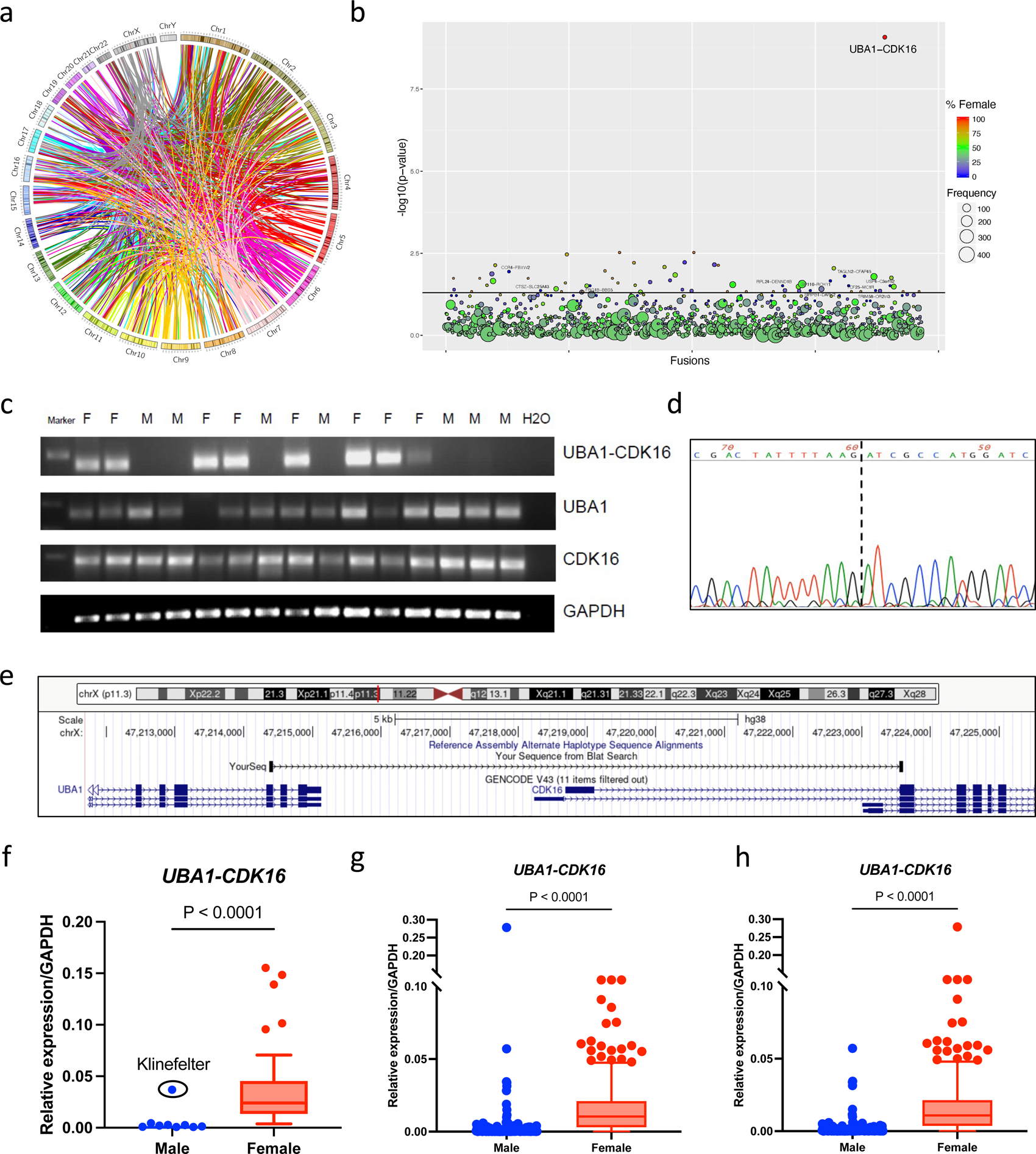
Discovery and validation of *UBA1-CDK16* chimeric RNA. (a) Circos plot depicted chimeric RNAs discovered across the genome from 426 paired-end RNA-seq data of GTEx whole blood. A total of 1,158 recurrent fusion events (>4) are shown, with “MM” fusions excluded. Lines denote the chimeric RNAs connecting two parental genes. (b) Bubble plot highlighted the significant sex-bias of *UBA1-CDK16* chimeric RNA. The size of each bubble indicates the frequency of candidate chimeric RNAs, while the color scale indicates the percentage of female samples that contain a particular chimera. The Y-axis plots the negative value of the log (base10) transformed p-value obtained via chi-squared tests. (c) RT-PCR validation of *UBA1-CDK16* chimeric RNA and parental genes in 15 whole blood samples from eight female (F) and seven male (M) healthy individuals. Primers flanking the predicted fusion junction site were used for PCR amplification from cDNA. Gel electrophoresis revealed the presence of the chimeric RNA in female samples. Primers specific for the parental genes were used to detect *UBA1* and *CDK16* transcripts, together with primers for the control gene *GAPDH* (Table S1). (d) A partial sequence of the RT-PCR product of the chimeric RNA obtained by Sanger sequencing. (e) Configuration of the Sanger sequenced part of chimeric RNA is depicted on the UCSC Genome Browser (hg38). (f) RT-qPCR analysis of the *UBA1-CDK16* chimeric transcript in 104 RNA samples obtained from GTEx, including 64 males and 40 females. Expression was normalized to that of *GAPDH*. (g) RT-qPCR analysis of *UBA1-CDK16* chimeric transcript in 1,252 clinical samples from UVa hospital, including 727 males and 525 females, revealed a significant difference between the sexes (p=5.6E-56). (h) Reanalysis of the data shown in (g) after correcting the sex of some of the outliers based on the presence or absence of the Y-linked gene, *DDX3Y*. Sex annotation of several samples was changed, resulting in 736 males and 516 females. The p-value is 2.7E-53.

With the aim to explore the role of chimeric RNAs in sexual dimorphism, we compared the distribution of selected chimeric RNAs between sexes. Notably, we discovered a chimeric transcript, *UBA1-CDK16,* which exhibited significant bias towards female expression (24/157 in females and 0/269 in males) (**Fig. 1b**). Recognizing the sensitivity limitations of RNA-sequencing and bioinformatic detection, we conducted further validation of this female-specific expression pattern through RT-PCR on RNA extracted from 15 whole blood samples (8 females and 7 males). PCR primers for *UBA1-CDK16* were designed at the predicted junction sites identified by EricScript, while primers for each wild-type parental genes were designed at sequences not involved in chimeric RNA (**Table S1**). Our experimental results confirmed the exclusive female expression pattern of *UBA1-CDK16.* Intriguingly, transcripts from the two parental genes did not exhibit any sex-biased expression (**Fig. 1c and S1a**). Furthermore, we conducted Sanger sequencing of the chimeric PCR product to confirm the junction sequence (**Fig. 1d),** which revealed the fusion of the last to the third exon (exon 24 based on RefSeq gene NM_003334) of *UBA1* with the second exon (exon 2 based on RefSeq gene NM_006201) of *CDK16* (**Fig. 1 e**). When analyzing RPKM (Reads Per Kilobase per Million mapped reads) of the parental genes from the GTEx whole blood dataset, which contains reads of the chimeric transcript, we found no statistically significant sex difference in *CDK16* expression, but a higher level of *UBA1* expression in females (**Fig. S1b**). Nonetheless, the parental genes are expressed in both sexes. The female specificity of the chimeric RNA suggests a specific mechanism contributing to this sex-biased chimeric transcript expression and a possible functional role independent of parental genes.

To further validate the pronounced female expression bias of *UBA1-CDK16*, we conducted RT-qPCR analysis on an additional 104 whole blood RNA samples directly obtained from GTEx (**Fig. S2a**). This expanded analysis revealed striking sex differences, as all female samples exhibited positive expression, while all male samples showed negative expression, except for a single outlier from a Klinefelter syndrome (XXY) donor (**Fig. 1f**). Our investigation found no correlation between the expression of *UBA1-CDK16* and factors such as age, BMI, or race (**Fig. S2b and c**). Furthermore, we observed no differential expression of *UBA1-CDK16* during different phases of menopause, indicating an independent regulation from sex hormones (**Fig. S2d**). In summary, our findings from both bioinformatic identification and experimental validation, have unveiled a novel *UBA1-CDK16* chimeric RNA that exhibits significant sex differences in expression among individuals with one (XY) or two X chromosomes (XX and XXY).

### Further validation of female specificity of *UBA1-CDK16* chimeric RNA in 1,252 clinical blood samples

To further confirm the sex-biased expression of the chimeric RNA, we took advantage of a large number of clinically discarded blood samples. Previous evaluation of the shelf-time effect on the stability of various transcripts supported the feasibility of using these leftover samples for RNA expression study. We then obtained 1,252 whole blood samples, comprising 516 females and 736 males (**Fig. S3a**), and extracted RNA from the buffy coat layer. In accordance with our previous findings, the majority of female samples exhibited positive *UBA1-CDK16* expression (471 out of 516), while a significant proportion of male samples remained below the detectable level (542 out of 736). The sex-biased difference in *UBA1-CDK16* expression was found to be highly significant (P = 5.6E-56) (**Fig. 1g**). Consistent with GTEx samples, our analysis revealed no statistically significant contributions of age, race, BMI, or menopause phases to the variation in *UBA1-CDK16* levels (**Fig. S3b, c, and d**).

However, our analysis did reveal the presence of outlier specimens. We identified 75 individuals classified as “females” who exhibited *UBA1-CDK16* chimeric RNA levels below the male average, while 8 individuals classified as “males” had levels higher than the female average. To address the possibility of specimen labeling errors and potential transgender situations, we examined the expression of a Y-linked gene, *DDX3Y*, in these outlier samples. Indeed, our analysis revealed that 12 of the initially classified “females” as biological males, and 3 of the initially classified “males” as biological females. The significant sex difference of *UBA1-CDK16* was further confirmed after Y-chromosome correction (**Fig. 1h**).

The presence of outlier samples in this clinical collection, but not in the health donors or GTEx donors may be related to the specific diseases’ prevalence among these patients. We then examined the clinical diagnostic codes for the outlier patients’ hospital visits. Even though we found associations of female outliers with chronic obstructive pulmonary disease, pulmonary collapse, and colostomy, and male outliers with upper respiratory infections and heart valve disorders, none of them reached statistical significance. Thus, a larger collection of outlier populations is needed.

### *UBA1-CDK16* is a product of cis-SAGe from the inactive X chromosome

In female cells, random inactivation of one of the X-chromosomes serves to compensate for the dosage balance with males. However, both *UBA1* and *CDK16* genes are known to escape X-inactivation^27^. Given the female-specific characteristics of *UBA1-CDK16*, we suspected that the chimeric RNA is produced from the inactive X chromosome. We then obtained clinical samples from several Klinefelter Syndrome patients and one Turner Syndrome patient. Consistent with our hypothesis, we observed positive expression of *UBA1-CDK16* in Klinefelter Syndrome patients with the XXY genotype, while it was absent in the Turner Syndrome patient, who possesses a single X genotype (**Fig. 2a**). This finding suggests that the *UBA1-CDK16* chimeric RNA is exclusively expressed from the inactive X chromosome.

**Figure 2.**
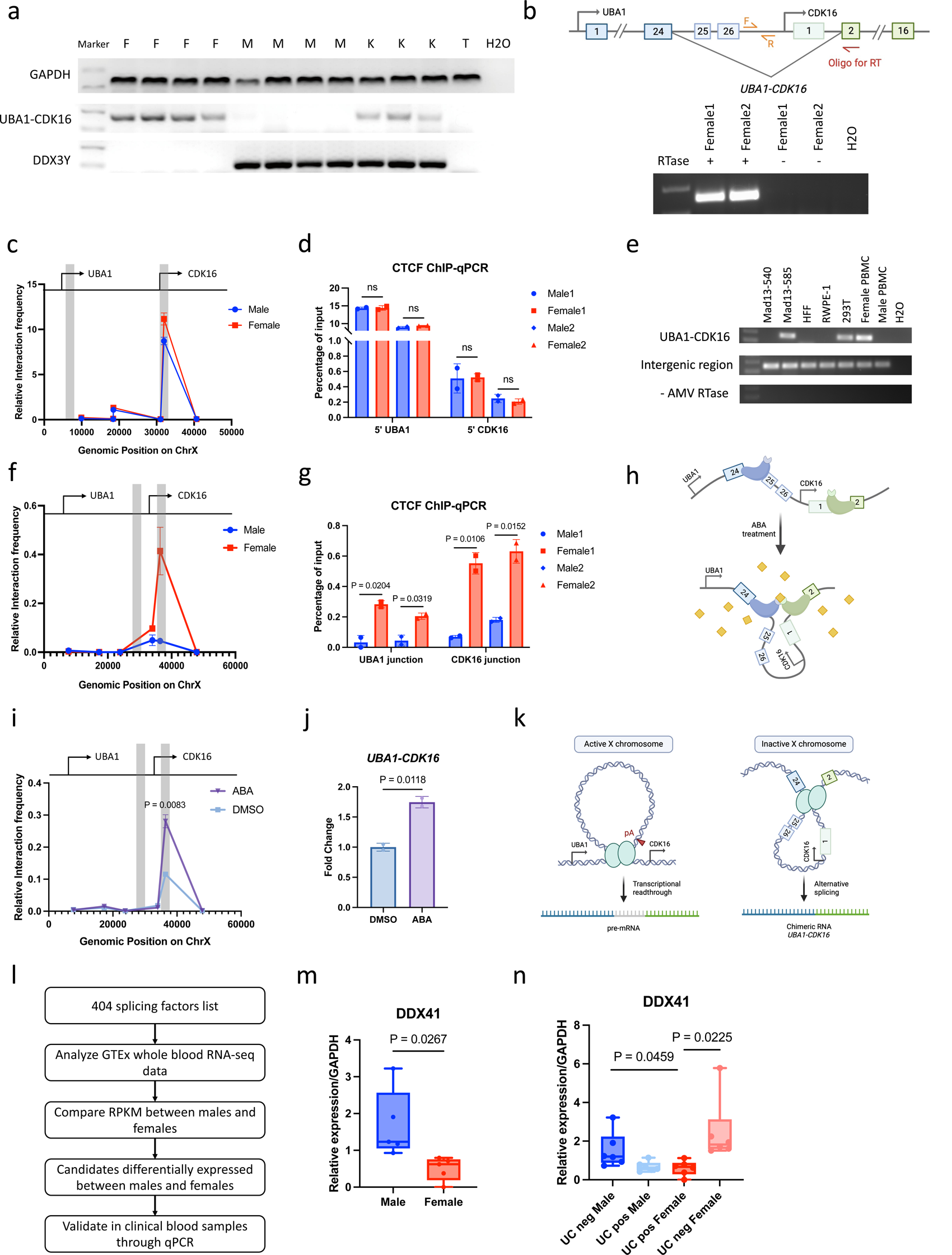
Mechanism of *UBA1-CDK16* chimeric RNA expression from the inactive X chromosome. (a) Gel electrophoresis revealed the RT-PCR positive detection of *UBA1-CDK16* in four females (F) and three Klinefelter syndrome patients (K) but absent in four males (M) and one Turner syndrome patients (T). Y chromosome was confirmed by the positive detection of *DDX3Y* in four males and three Klinefelter syndrome patients. *GAPDH* was used as control. (b) Schematic of the cis-SAGe PCR shows the oligo for reverse transcription annealed to the exon2 of *CDK16* and the PCR primers flanking the intergenic region. Gel electrophoresis showed cis-SAGe PCR products from two female PBMCs treated with DNaseI. Samples without treatment by RT enzyme were used as negative control. (c) 3C assay measuring *UBA1-CDK16* locus-wide crosslinking frequencies in male and female PBMCs. Samples were fragmented by BamHI. The 5’ *UBA1* fragment was used as the anchor. Its interaction frequency with other 5’ *CDK16* fragment was highlighted in grey. (d) ChIP-qPCR demonstrated CTCF-binding at the 5’ *UBA1* and 5’ *CDK16* loci in two pairs of male and female PBMCs. Two-tailed unpaired t-tests revealed no sex differences in CTCF-binding on both sites. (e) RT-PCR revealed the mature *UBA1-CDK16* chimeric transcript expression in female cell lines Mad13-585, 293T, and female PBMCs. Cis-SAGe PCR revealed the intergenic region detection in all cell lines. Samples without treatment by RT enzyme were used as negative control. (f) 3C assay measuring *UBA1-CDK16* locus-wide crosslinking frequencies in male and female PBMCs. Samples were fragmented by BglII. The *UBA1* junction fragment was used as the anchor. Its interaction frequency with other *CDK16* junction fragment was highlighted in grey. (g) ChIP-qPCR demonstrated CTCF-binding at the *UBA1* and *CDK16* junction loci in two pairs of male and female PBMCs. Two-tailed unpaired t-tests revealed significant higher frequencies of CTCF-binding on both sites in females. (h) Schematic of CLOuD9 with sgRNAs targeting junction sites. Addition of abscisic acid (ABA, yellow) brings two complementary CLOuD9 constructs (blue and green) into proximity, inducing chromatin loop formed between junction sites. (i) 3C assay measuring *UBA1-CDK16* locus-wide crosslinking frequencies in HEK293T cells after 3 days of treatment with ABA. CLOuD9 construct target regions were highlighted in grey with *UBA1* junction fragment as the anchor. An increased interaction between target junction regions was observed. (j) CLOuD9-induced chromatin looping at the junction sites resulted in induction of *UBA1-CDK16* expression conducted by RT-qPCR. (k) Summary schematic of chromatin structure at *UBA1-CDK16* locus on active and inactive X chromosome. (l) Pipeline of discovering sex-differential expressed splicing factors from GTEx whole blood RNA-seq data. (m) Validation of *DDX41* expression by RT-qPCR in RNA extracted from five male and female buffy coats. (n) RT-qPCR detection of *DDX41* expression in outlier clinical blood samples from Fig. 1h.

*UBA1* and *CDK16* are neighboring genes located on the X chromosome. To investigate the mechanisms responsible for the formation of this chimeric RNA, we initially examine whole genome sequencing data from GTEx and confirmed that it is not a product of an interstitial deletion (**Fig. S4a**). *UBA1* and *CDK16* transcribed in the same orientation, fitting the characteristics of cis-splicing between adjacent genes (cis-SAGe). To further confirm that *UBA1-CDK16* is a product of cis-SAGe, we designed a cis-SAGe RT-PCR assay to detect the precursor readthrough mRNA. In this assay, we used an oligo annealing to exon2 of *CDK16* to specifically reverse transcribe the precursor readthrough mRNA. The readthrough precursor was then successfully detected using intergenic primers in female PBMCs (**Fig. 2b**). To ensure the elimination of DNA contamination, RNA was treated with DNaseI, and the no RT enzyme control indicated the complete removal of DNA contaminants.

### A 5’-5’ chromatin loop generates sex-independent transcriptional readthrough

To delve further into the molecular mechanisms underlying the formation of *UBA1-CDK16* through cis-SAGe, we examined the Xi chromatin structure by comparing male and female peripheral blood mononuclear cells (PBMCs). Prior studies have reported that the intergenic chromatin loops formed between promoters can promote the transcriptional readthrough by obstructing the poly-A recognition site^28^. To investigate this, we conducted Chromosome Confirmation Capture (3C), which revealed a prominent chromatin loop between the 5’ regions of *UBA1* and *CDK16*, characterized by notable high interaction frequencies (**Fig. 2c**). Subsequently, we performed CTCF Chromatin Immunoprecipitation (ChIP) analysis and identified a significant CTCF binding frequency at the predicted CTCF-binding sites around the loop anchor sites (**Fig. S4b and c**). This notable CTCF binding at both 5’ regions of *UBA1* and *CDK16* was confirmed in two pairs of *PBMCs*, providing further evidence of the anchor sites for the observed chromatin loop (**Fig. 2d**). Notably, no sex differences were observed in either chromatin structure or CTCF binding frequency.

To further investigate the role of this 5’-5’ chromatin loop in *UBA1-CDK16* expression, we induced this chromatin loop in HEK293T cells through CRISPR-mediated chromatin loop organized using dCas9 (CLOuD9)^29^. HEK293T, known for its high transfection efficiency, is a female cell line with low *UBA1-CDK16* expression. Employing sgRNAs targeting the identified loop anchor sites, we induced an *in vitro* chromatin loop through dCas9 dimerization, followed by treatment with abscisic acid (ABA) (**Fig. S4d**). Facilitated by this chromatin loop, we observed a significant increase of transcriptional readthrough precursors, as detected by cis-SAGe RT-PCR. Importantly, this increase in readthrough precursors did not impact the mature *UBA1-CDK16* expression (**Fig. S4e**).

Given that the chromatin loop between 5’-5’ regions of *UBA1* and *CDK16* is detected in both female and male samples, we postulated that the transcriptional readthrough precursors present in both males and females. To test this hypothesis, we measured *UBA1-CDK16* and its precursor in seven different cell lines, encompassing both male and female cell lines. Remarkably, we observed the presence of readthrough precursors in all cell lines, irrespective of gender (**Fig. 2e**). However, the mature chimeric transcript *UBA1-CDK16* was specifically produced in female cell lines. These findings collectively highlight the sex-independent role of the 5’-5’ chromatin loop in mediating transcriptional readthrough and suggest a potential female-specific alternative splicing event leading to the expression of *UBA1-CDK16*.

### A female-specific chromatin loop is formed between*UBA1-CDK16* junction sites

It has been reported that CTCF-mediated chromatin loops can regulate alternative splicing ^30^. Considering the distinct female-specific expression characteristics of *UBA1-CDK16* and the presence of its readthrough precursors in both sexes, we posited that a specific chromatin structure on the Xi chromosome might contribute to the formation of *UBA1-CDK16*. To test this hypothesis, we compared chromatin interactions between the junction sites in male and female PBMCs using 3C. Our study identified that a chromatin loop between the junction sites with a higher level of interaction in female PBMCs (**Fig. 2f**). Notably, we also detected CTCF binding around the anchor sites (**Fig. S4f and g**), and in two pairs of donors, we observed significant female-specific CTCF binding at both junction sites (**Fig. 2g**).

To prove the role of this junction chromatin loop in *UBA1-CDK16* expression, we conducted CLOuD9 experiments with sgRNAs targeting the identified junction anchor sites (**Fig. 2h**). This manipulation induced a significant chromatin interaction between junction sites (**Fig. 2i**) and resulted in an increase of *UBA1-CDK16* expression (**Fig. 2j**). Therefore, this female-specific chromatin loop, responsible for the proximity of junction splicing sites, promotes the expression of the chimeric transcript *UBA1-CDK16*.

In summary, our findings highlight that while a chromatin loop formed between the 5’ regions of *UBA1* and *CDK16* mediates transcriptional readthrough in both males and females, the female-specific *UBA1-CDK16* mature chimeric transcript is facilitated through a specific intergenic chromatin loop formed between junction sites on the inactive X chromosome (**Fig. 2k**).

### Sex-differential splicing factors correlated with*UBA1-CDK16* expression

To uncover the splicing factors involved in *UBA1-CDK16* expression, we first aimed to identify splicing factors with sex-differential expression. We obtained a list of 404 splicing factors from a study on the landscape of splicing factors^31^. We then analyzed their RPKM values from GTEx whole blood RNA-seq data and compared them between sexes (**Fig. 2l**). This analysis revealed 12 splicing factors with significant sex differences in whole blood (**Fig. S5a**). We performed RT-PCR validation using RNA samples extracted from the buffy coat layer of clinical blood samples. Among the candidates, six splicing factors were found to be differentially expressed (**Fig. S5b**), with only *DDX41* showing consistent differential expression with GTEx sequencing analysis, exhibiting significantly higher expression in males (**Fig. 2m**). The discrepancy may be due to the difference between sample types, with RT-PCR utilizing buffy coat, while GTEx sequencing analysis involves whole blood samples.

Taking the advantage of the outlier clinical samples with abnormal expression of *UBA1-CDK16*, we aimed to investigate the correlation between candidate splicing factors and *UBA1-CDK16* expression. We examined the expression of these splicing factors in males, females, “males” positive for *UBA1-CDK16,* and “females” negative for *UBA1-CDK16.* The significant higher expression of *DDX41* in female outliers provided additional support for its potential regulatory roles in modulating *UBA1-CDK16* expression (**Fig. 2n**). Other splicing factors, including *ZRSR2*, SRPK1, *ALYREF*, and *YBX1*, also exhibited differential expression in outliers (**Fig. S6**). The precise mechanism underlying *UBA1-CDK16* splicing events, potentially involving multiple factors, requires further investigation.

### Existence of *UBA1-CDK16* during evolution

Based on GTEx data analysis, minimal expression of *UBA1-CDK16* was observed in tissues other than blood. To confirm the enrichment of *UBA1-CDK16* in blood, we experimentally measured its expression in various female tissues including adrenal gland, bladder, liver, lung, skeletal muscle, small intestine, stomach, and tonsil. While low levels of *UBA1-CDK16* were observed in tonsil and liver, we confirmed its significant enrichment in the buffy coat (**Fig. S7a**).

To investigate the functional significance of the chimeric transcript *UBA1-CDK16* in blood and explore its presence in an evolutionary context, we examined its expression across species from *Mus musculus* to *Homo Sapiens* (**Fig. 3a**). Our findings unveiled the presence of *Uba1-Cdk16* in both male and female mice (**Fig. 3b**). However, Sanger sequencing of the chimeric PCR product indicated the existence of a distinct e25e2 isoform (exon25 of Uba1 and exon2 of Cdk16 based on the Refseq gene NM_009457 and NM_011049), different from the human e24e2 form (**Fig. 3c**). As we traced its evolutionary trajectory in non-human primates, we observed a notable gain of the e24e2 isoform in marmosets, but without a sex-biased expression pattern for either form (**Fig. 3d**). In baboons, the e25e2 isoform further evolved to become female-specific, coexisting with e24e2 isoform in both sexes (**Fig. 3e**). In rhesus monkeys, both e24e2 and e25e2 isoforms exhibited female-specific expression (**Fig. 3f**). The presence of two isoforms and e25e2 being female specific in baboons, while both isoforms specific to female in rhesus monkeys suggest two routes of evolution converge in the end result of e24e2 being the exclusive isoform present in the human females. The gain of the female-specific chimeric RNA *UBA1-CDK16* during dynamic evolution strongly supports its potential significant functional roles in the blood system.

**Figure 3.**
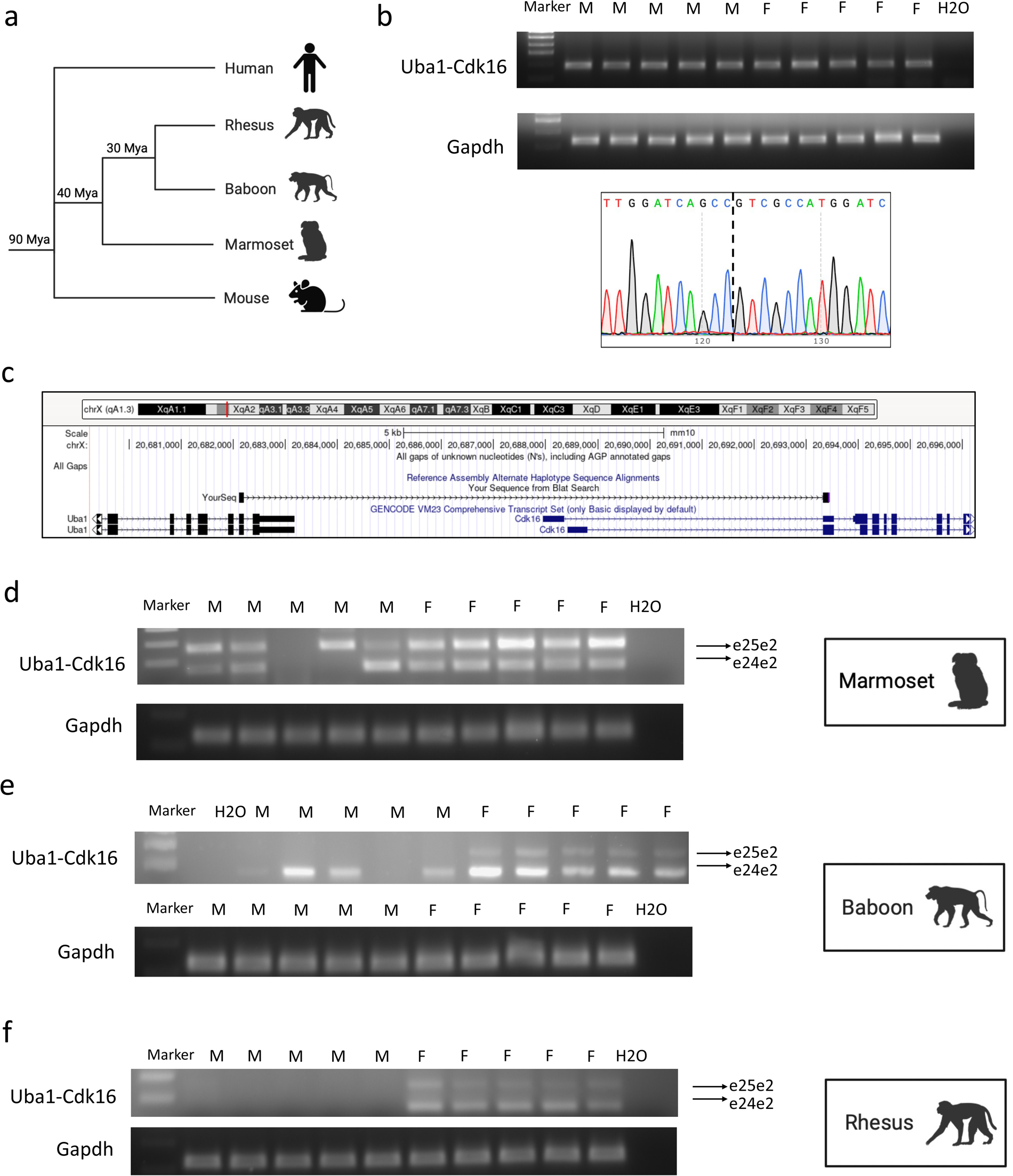
Comparative study of *UBA1-CDK16* expression during evolution. (a) The schematic phylogenies represent evolutionary relationships among mice, non-human primates, and humans. (b) Gel electrophoresis revealed the presence of the chimeric RNA *Uba1-Cdk16* in both male and female mice. RNA was extracted from buffy coat. A partial sequence of the RT-PCR product of the chimeric RNA obtained by Sanger sequencing is shown below. (c) Configuration of the Sanger sequenced part of chimeric RNA is viewed on the UCSC Genome Browser (mm10). (d,e,f) Gel electrophoresis revealed the presence of different isoforms of the chimeric RNA *Uba1-Cdk16* in marmosets, baboons, and rhesus monkeys.

### *UBA1-CDK16* inhibits CD11b expression during myeloid differentiation

Blood, as a vital line of body’s defense, plays a critical role in immune system. To investigate the potential function of *UBA1-CDK16* in blood immune system, we sorted specific blood cell subsets from peripheral blood and examined the expression of *UBA1-CDK16*. Our findings revealed a significant enrichment of *UBA1-CDK16* in myeloid lineage cells, including red blood cells and CD11b+ myeloid cells, in comparison to its expression in lymphoid lineage cells including NK cells, T cells, and B cells (**Fig. 4a**). To gain deeper insight into the dynamic expression of *UBA1-CDK16* within the myeloid lineage, we initially investigated its presence in hematopoietic stem cells (HSCs). Surprisingly, this chimeric RNA is undetectable in CD34+ HSCs isolated from peripheral blood. However, both parental transcripts, *UBA1* and *CDK16,* were expressed in both CD34+ and CD34-cells at similar levels (**Fig. S7b**). We then induced the differentiation of HSCs into the myeloid lineage by treating them with a combination of SCF, Flt3L, IL-3, IL-6, and GM-CSF. Our data revealed a progressive increase in *UBA1-CDK16* expression on Day 3 and Day 5 during the myeloid differentiation (**Fig. 4b**).

**Figure 4.**
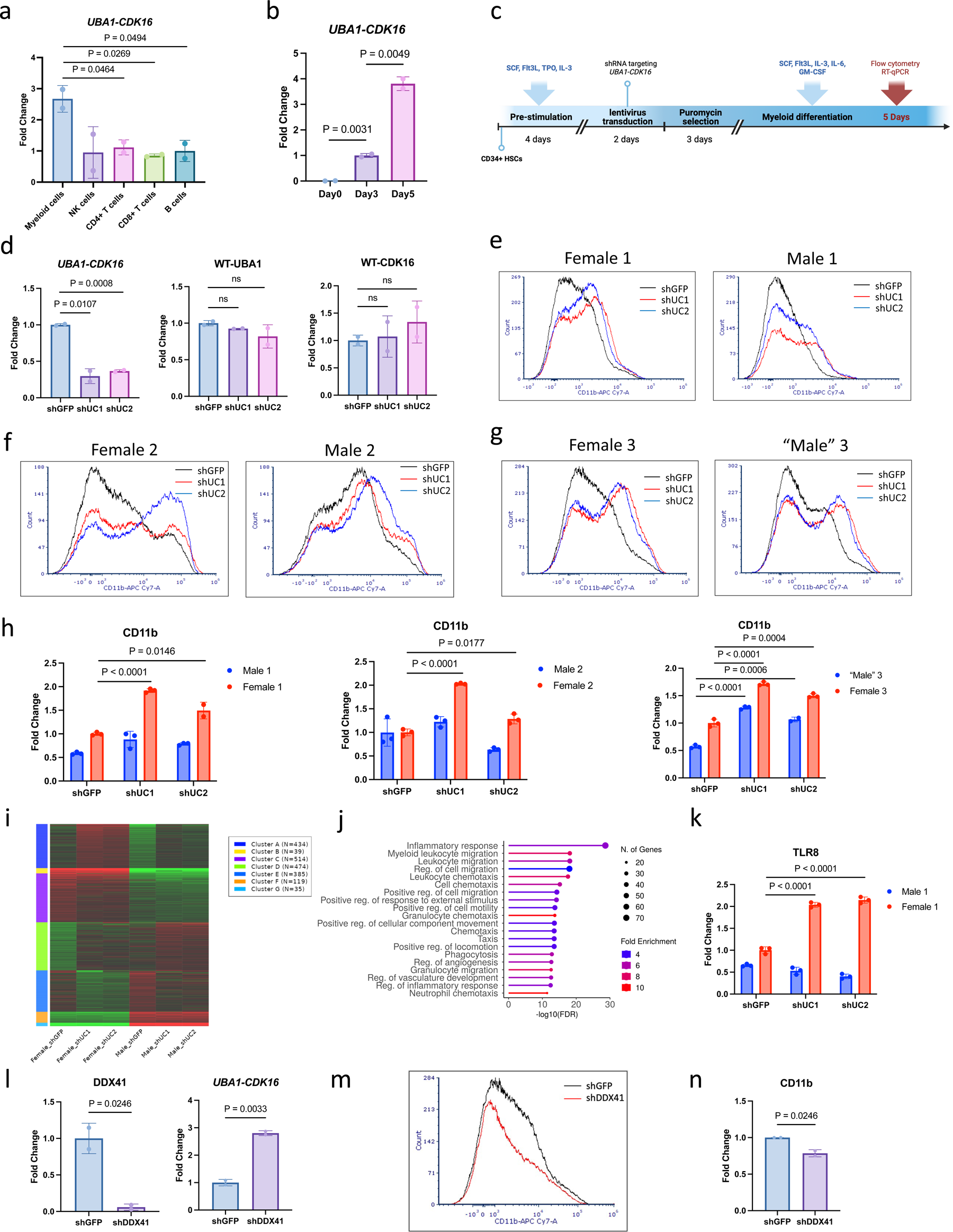
Function of *UBA1-CDK16* during myeloid differentiation. (a) RT-qPCR detection of chimeric RNA *UBA1-CDK16* in myeloid cells, NK cells, CD4+ T cells, CD8+ T cells, and B cells sorted from human PBMCs. (b) RT-qPCR detection of chimeric RNA *UBA1-CDK16* on Day 0, 3, and 5 of CD34+ cells myeloid differentiation. (c) Experimental timeline for shRNA transduction of expanded CD34+ cells followed by myeloid differentiation. (d) The effect of *UBA1-CDK16* knockdown by shUC1 and shUC2. Parental genes expression was not affected by shUC. shGFP was used as negative control. (e,f,g) The fluorescent intensity distribution of CD11b was analyzed in three pairs of CD34+ cells donors. Gated out single cells were used for FACS analysis. (h) CD11b mRNA expression detected by RT-qPCR. RNA samples were extracted from day 5 of myeloid differentiated cells. (i) Hierarchical clustering indicated the substantial differences in genes induced by *UBA1-CDK16* knockdown in males and females. (j) Enriched Gene Ontology (GO) terms for differentially expressed genes in cluster A. Enriched biological processes were listed based on false discovery rate (FDR) ranking. Dot size indicates the number of genes, while the color scale indicates fold enrichment. (k) RT-qPCR validation of *TLR8* in the same RNA samples used for sequencing. The relative expression level was normalized against *GAPDH*, and fold change was normalized against female shGFP group. (l) The effect of *DDX41* knockdown on the upregulation of *UBA1-CDK16* during myeloid differentiation. (m) The fluorescent intensity distribution of CD11b in shDDX41 vs. shGFP. (n) RT-qPCR indicated a decrease of CD11b mRNA level upon DDX41 knockdown.

Recognizing the significant upregulation of *UBA1-CDK16* during this timeline, we decided to study its function during *in vitro* myeloid differentiation. We designed two shRNAs targeting the chimeric junction sequence to reduce *UBA1-CDK16* expression specifically in expanded HSCs through lentivirus transduction (**Fig. 4c**). Three days after initiating myeloid differentiation, we observed an efficient knockdown of *UBA1-CDK16*, with no changes in the expression of the parental transcripts (**Fig. 4d**). Following five days of differentiation, our flow cytometry analysis revealed the prevalence of a myeloblast subset characterized by the CD33+CD11b+ population (**Fig. S7c**). Upon *UBA1-CDK16* knockdown, we observed a notable increase in the CD11b+ population, as indicated by enhanced fluorescence intensity in the female donor (**Fig. 4e**). In contrast, the male donor, utilized as a shRNA off-target control, exhibited a CD11b fluorescence distribution similar to that of the shGFP control (**Fig. 4e**). To eliminate the possibility of donor-related variability, we replicated this experiment using two additional pairs of HSC donors. In the second pair, we consistently observed an increase in the CD11b+ population resulting from the loss of *UBA1-CDK16* in a female specific manner (**Fig. 4f**). In the third pair, intriguingly we observed the same phenotype in both the female and “male” donors, with significant shift in the fluorescence intensity of CD11b with both shRNAs (**Fig. 4g**). We eventually found out that this male donor was positive for *UBA1-*CDK16, and the shRNAs were also effective silencing the chimera in this aberrant male donor. We then conducted RT-qPCR analysis and consistently detected an elevation in CD11b mRNA levels in all *UBA1-CDK16* positive donors, including the aberrant “male” donor (**Fig. 4h**). These findings strongly suggest a crucial role for *UBA1-CDK16* as a regulator of hematopoiesis, potentially functioning as a checkpoint for excess myeloid differentiation.

### *UBA1-CDK16* is involved in inflammatory response pathways

To explore the potential downstream target of *UBA1-CDK16* involved in the regulation of myeloid differentiation, we analyzed the RNA-seq data obtained from *UBA1-CDK16* shRNA transfected samples of both female and male. Utilizing K-means clustering^32^ (**Fig. S8a**), we identified a distinct cluster of genes that exhibited differential expression specific to the female knockdown samples (**Fig. 4i**). By performing the Gene Ontology analysis, this cluster of genes were found to regulate immune and inflammatory responses (**Fig. 4j**). To validate the potential downstream targets, we conducted RT-qPCR analysis on samples from all three pairs of donors. Our results demonstrated a significant upregulation of *TLR8* in response to the loss of *UBA1-CDK16* including in the aberrant male expressing *UBA1-CDK16* (**Fig. 4k and S8b**). To further investigate the pathways *UBA1-CDK16* may be involved in inflammatory responses, we analyzed the hallmark pathways correlated with the identified cluster of genes. TNF-α signaling via NF-κB pathway was found to be significantly enriched (**Fig. S8c**). Experimental validation results confirmed that *IL1B* and *LRP1* involved in NF-κB signaling pathway were upregulated upon *UBA1-CDK16* knockdown in females (**Fig. S8d**). Our findings strongly suggest that this female-specific chimeric transcript *UBA1-CDK16* plays a pivotal role in mediating inflammatory responses by regulating the TNF-α/NF-κB signaling.

### *DDX41* may promote myeloid differentiation by negatively regulating*UBA1-CDK16*

As mentioned above, we identified a splicing factor *DDX41,* which displayed a negative correlation with *UBA1-CDK16* expression. To test the impact of *DDX41* during *in vitro* myeloid differentiation, we employed shRNAs to knock down *DDX41* in female HSCs and subsequently initiated the myeloid differentiation. Remarkably, the decrease of *DDX41* resulted in a significant increase in *UBA1-CDK16* expression (**Fig. 4l**). Notably, the elevated levels of this myeloid checkpoint inhibitor were accompanied by a decrease in the CD11b+ population, as indicated by reduced CD11b fluorescence intensity (**Fig. 4m and S7d**). Consistently, the mRNA levels of CD11b showed a decrease upon *DDX41* knockdown (**Fig. 4n**).

### Abnormal expression of *UBA1-CDK16* in COVID-19 patients

During the recent COVID-19 pandemic, multiple studies have highlighted a gender disparity, with males experiencing more severe COVID-19 symptoms and higher mortality rates compared to females. These differences have been linked to sex-biased immune responses, wherein females exhibit a more robust T cell activation, while males demonstrate stronger innate immune responses^33^. Given our understanding of *UBA1-CDK16*’s potential role in regulating myeloid differentiation, we aimed to further investigate its involvement in responding to COVID-19 infection. We examined the expression levels of *UBA1-CDK16* in female COVID-19 patients categorized by symptom severity, including asymptomatic, mild, severe, and critically ill or death. We used whole blood samples from healthy individuals as control, where *UBA1-CDK16* was consistently present in all females and absent in males. Our findings showed that *UBA1-CDK16* was normally expressed in all 22 asymptomatic female COVID-19 patients (**Fig. 5a**). However, as the severity of COVID-19 symptoms increased, a subgroup of female patients lost the expression of *UBA1-CDK16* (**Fig. 5a**). While quantifying the level of *UBA1-CDK16* expression, we observed a dramatic decrease in chimeric RNA levels, which correlated with the increasing severity of symptoms (**Fig. 5b**). To further investigate the *UBA1-CDK16* correlated sex-biased COVID-19 phenotypes, we compared blood test results between female patients who tested positive and negative for *UBA1-CDK16* in each symptom group (**Fig. S9a, b**, and **c**). We noticed a notably higher level of neutrophil counts in mild female patients who had lost *UBA1-CDK16* expression (**Fig. 5c**). However, no obvious difference was observed in lymphocyte counts (**Fig. 5d**). Recent studies have identified the Neutrophils to Lymphocytes Ratio (NLR) as an independent COVID-19 prognosis marker^34,35^. Consistently, we observed a significant elevation of NLR in the symptomatic patients who tested negative for *UBA1-CDK16* (**Fig. 5e**). These findings provided additional evidence supporting the inhibitory regulatory role of *UBA1-CDK16* in myeloid cell differentiation, and its potential implications in the severity of COVID-19.

**Figure 5.**
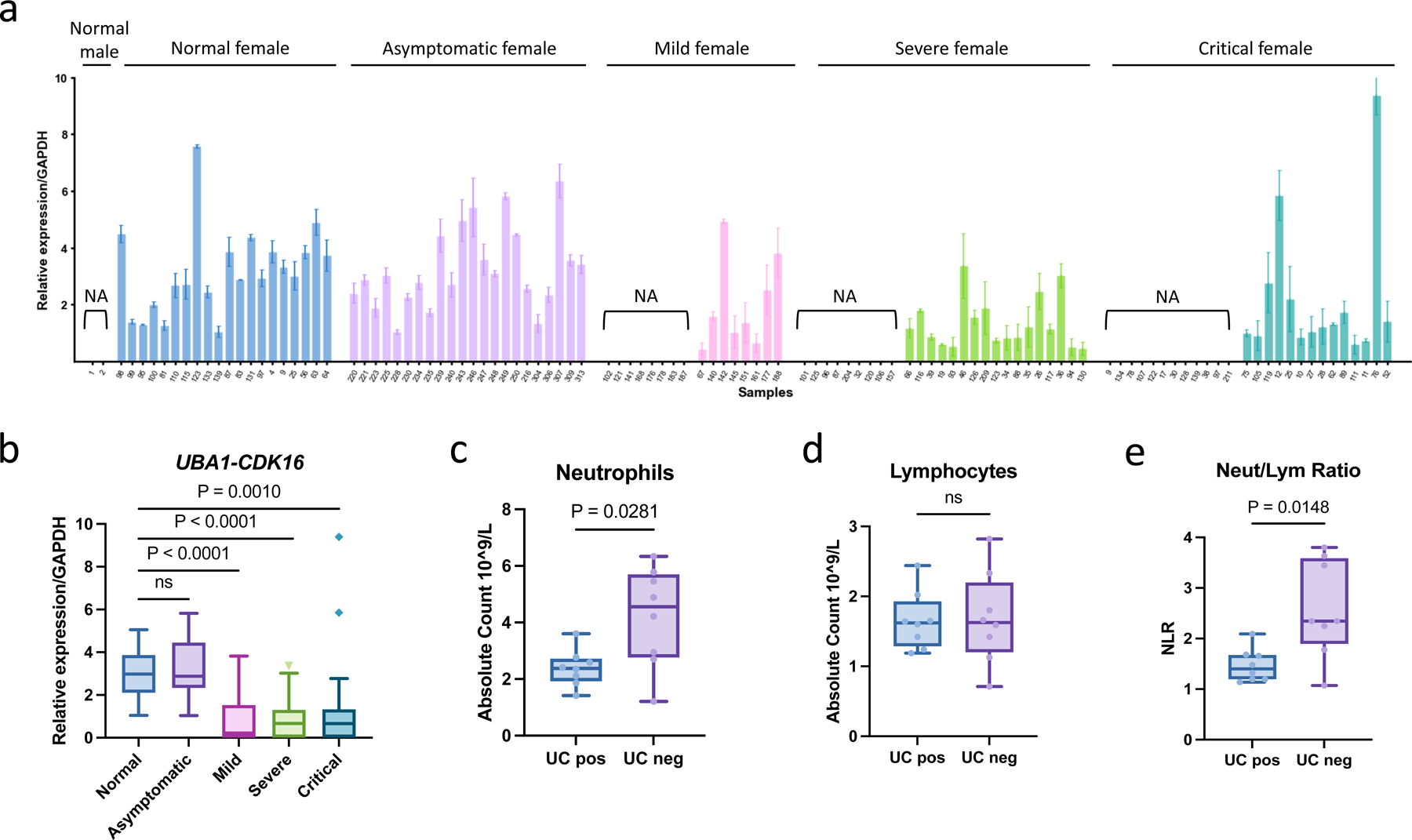
Abnormal expression of *UBA1-CDK16* in COVID-19 female patients. (a) RT-qPCR revealed the expression of *UBA1-CDK16* in each female COVID-19 patient. Normal females detected positive for *UBA1-CDK16* were used as positive control, while normal males were used as negative control. (b) RT-qPCR revealed the expression of *UBA1-CDK16* in female COVID-19 patients grouped by symptoms, including asymptomatic, mild, severe, and critical cases. (c,d,e) Neutrophil and lymphocyte counts and the neutrophil-to-lymphocyte ratio (NLR) were evaluated in patients who tested positive or negative for *UBA1-CDK16*.

### Other chimeric RNAs expressed from escaping zone of X inactivation are not female-biased

We postulated that transcriptional readthrough from Xi can produce other female-specific chimeric RNAs in addition to *UBA1-CDK16*. We previously reported several common features about the parental genes of cis-SAGe chimeric RNAs: (1) the gene pairs consistent of adjacent genes transcribed in the same orientation; (2) the parental genes tend to be close to each other (<30kb) when compared to other gene pairs within the genome; and (3) they are usually multi-exonic, and 4) the chimera tends to involve specific exons of the parental genes^36^. Consistently, *UBA1* and *CDK16* are only 3kb apart, and the chimeric RNA involves the third-to-last exon of *UBA1* joining to the second exon of *CDK16*. We then examined the regions containing genes that escape X-inactivation^27^. Using the parameters listed above, we identified two additional gene pairs that may produce female-biased chimeric RNAs, including: *SYAP1* with *TXLNG* (24kb apart), and *TCEANC* with *RAB9A* (25kb apart). According to the common exon combinations of cis-SAGe fusion, we designed a forward primer annealing to the second-to-last, or the third-to-last exon of the 5’ gene, and a reverse primer annealing to the second or third exon of the 3’ gene. Indeed, we were able to identify and validate the presence of two chimeric RNAs, *SYAP1-TXLNG* and *TCEANC-RAB9A* (**Fig. S10a and b**). However, expression analyses of these two chimeric RNAs in the same set of 15 whole blood samples used to examine *UBA1-CDK16* chimeric RNA, detected only a few positive samples, with no evidence of a female bias (**Fig. S10c**), further suggesting that additional mechanism in addition to transcriptional readthrough contributes to the sex specificity of *UB1-CDK16*.

## Discussion

Chimeric RNAs were once thought to be exclusively generated by gene fusion at the DNA level, thus unique to cancer cells^6,8^. Recent studies have demonstrated that these RNAs can also result from intergenic splicing and be part of normal physiology^11,37,38^. In this study, we uncovered a novel sex-specific chimeric RNA, *UBA1-CDK16,* almost exclusively found in blood. This chimeric RNA is the result of cis-SAGe, occurring solely from the inactive X chromosome. As a result, it is expressed in females, while the parental genes are expressed in both males and females. This discovery supports the notion that chimeric RNAs may be a means to expand the transcriptome and diversify the pool of encoded genetic information.

The identification of chimeric RNAs can vary depending on the methods employed. Several software packages have been developed for identifying chimeric RNAs from RNA-seq data. In our study, we opted to use EricScript based on prior comparative research^25,39^. However, it is important to notice that the frequency of chimeric RNA detection is inherently constrained by the bioinformatics tools utilized. We therefore conducted a comprehensive experimental validation. Experimentally, GTEx blood samples exhibited a 100% detection rate in females, confirming the female-specific characteristics of *UBA1-CDK16.* While in the clinical samples from hospital, the detection rate was approximately 92%. This discrepancy suggests the potential for abnormal *UBA1-CDK16* expression in specific disease contexts.

To emphasize the significance of the *UBA1-CDK16* chimeric transcript, it is essential to delve deeper into its unique expression pattern. In humans, both *UBA1* and *CDK16* escape X inactivation^27^, which triggers transcriptional readthrough initiated by the *UBA1* promoter. We have confirmed the previous findings regarding the mechanisms of transcriptional readthrough facilitated by chromatin interactions within the 5’ regions of parental genes^28^. However, it’s worth noting that this chromatin loop alone is insufficient for cis-SAGe. Our study highlighted the involvement of alternative splicing in cis-SAGe, a process mediated by differences in chromatin structure on the inactive X chromosome^40^. The distinct chromatin loop formed between the junction splicing sites distinguishes the chromatin structure between the active (Xa) and inactive (Xi) chromosomes, ultimately leading to the formation of the chimeric RNA in females. While we have established the involvement of *DDX41* in *UBA1-CDK16* expression, the roles of other splicing factors, potential histone modifications, and additional molecular mechanisms necessitate further investigation.

The evolutionary trajectory of *UBA1-CDK16* across species provides compelling evidence for its functional significance. While we observed its presence in a different isoform in both male and female mice, the emergence of a female-specific isoform in baboons and rhesus monkeys suggests that this chimeric RNA has evolved to play specialized roles. This underscores the importance of further investigating its functions and implications within the evolutionary context of the blood immune system and sexual dimorphism.

The functional study of *UBA1-CDK16* was largely limited by its exclusive expression in primary cells. We observed that the chimeric RNA was expressed at an undetectable level in lymphoblastoid cell lines, potentially attributed to EBV transformation or *in vitro* culture conditions. Therefore, conventional cell lines proved unsuitable for investigating the functions of *UBA1-CDK16.* To overcome this limitation, we evaluated the chimeric RNA expression levels during myeloid differentiation from primary CD34+ cells and observed a significant increase in *UBA1-CDK16* expression. We thus employed this experimental system to study the phenotypes effects influenced by *UBA1-CDK16* during myeloid differentiation.

Sexual dimorphism in hematological systems has been a subject of increasing interest, with evidence pointing towards the involvement of X-linked genetic factors. The X-chromosome inactivation (XCI) escape is likely to contribute to female-specific functions in immune cells^41,42^. A recent study has highlighted the role of Xist ribonucleoproteins in promoting female sex-biased autoimmunity^43^. Our findings here address the other side, and illustrate that females have some protective mechanism against excessive autoimmunity. Previous studies have reported that the higher dosage of TLR8 due to XCI escaping plays a major role in inflammatory responses and contributes to the high incidence of autoimmune diseases in females^42,44^. Our findings strongly suggest that this female-specific chimeric transcript, *UBA1-CDK16,* may play a pivotal role in mediating inflammatory responses by regulating the expression of TLR8. Upon knockdown of *UBA1-CDK16* in HSCs, followed by myeloid differentiation, we observed an elevated level of TLR8 expression and an increased population of myeloblasts, as indicated by the higher expression of CD11b. Although further investigations are needed to fully elucidate the precise mechanisms involved, our study has provided a significant step in uncovering the role of sexual transcriptomic variants in contributing to sex-biased immunity.

Appreciating the function of *UBA1-CDK16* in myeloid differentiation, we aimed to explore its potential implications in sex-biased diseases. Multiple studies have reported sex differences in the immune responses of COVID-19 patients^33,45^. In our study, we observed higher counts of neutrophils in mild COVID-19 patients who tested negative for *UBA1-CDK16*, further confirming the inhibitory role of this chimeric RNA in myeloid differentiation. While a subset of patients in severe and critical groups also tested negative for *UBA1-CDK16,* no difference in neutrophil counts was observed. It is possible that different stages of virus infection may involve additional immune regulatory mechanisms in addition to *UBA1-CDK16.* The increased Neutrophil-to-Lymphocyte Ratio (NLR), which serves as a prognosis marker^34,35^, observed in mild COVID-19 cases raises intriguing questions about the functional consequences of *UBA1-CDK16* loss in myeloid cells during infection. Negative detection of *UBA1-CDK16* may indicate poor outcomes for COVID-19 patients. Moreover, many immune-related disorders, such as cancer, infectious diseases, or autoimmune diseases, exhibit sex differences, especially in myeloid cells. In the future, we aim to screen for the potential abnormal expression of *UBA1-CDK16* in various sex-biased diseases. Given its enrichment in blood, this novel discovery may shed like on the potential use of this sex-specific chimeric RNA as a diagnostic or prognostic marker.

## Acknowledgements

High-performance computing systems and services were provided by the Data Science Institute and the other Computation and Data Resource Exchange (CADRE) partner organizations at the University of Virginia.

Flow cytometry data for this manuscript were generated in the University of Virginia Flow Cytometry Core Facility (RRid:SCR_017829) and is partially supported by the NCI Grant (P30-CA044579). Cell sorting was performed on the Cytek Aurora CS funded through the NIH S10 instrumentation grant (1S10OD034355-01).

## Author contributions

H.L. and X.S. conceived the idea, interpreted the data and wrote the manuscript; S.S., S.K., R.C., and J.E. performed bioinformatic analysis; L.F., C.L., F.Q., and A.L. performed experimental validation; X.S. conducted molecular experiments and functional studies; T.B. performed cell sorting; S.L and P.W. analyzed the COVID patients’ data. All authors read the final version of the manuscript and agreed to its publication.

## Materials and methods

### Bioinformatics Analyses

Raw RNA-seq datasets (7,059) for 44 different human tissues were downloaded from the Genotype-Tissue Expression (GTEx) Project and subjected to the Next Generation Sequencing Quality Control toolkit^46^ (http://www.nipgr.res.in/ngsqctoolkit.html) to filter out low-quality reads. Paired-end sequencing reads were mapped to the Human genome (version hg38) and analyzed using the software tool EricScript^25^ to identify candidate chimeric RNAs. Fusions with an Ericscore of less than 0.6 were filtered out. The occurrence and frequencies of candidate chimeric RNAs were then correlated with the sex of the donors. Pearson’s chi-squared tests were performed between observed fusion frequencies and expected sample frequencies in male and female samples to generate a bubble plot. In this plot, data points are replaced with bubbles and the Y-axis plots the negative value of the log(base10) transformed p-value obtained via chi-squared tests, with the size of each bubble representing the total frequency of each chimera in GTEx blood samples, and the color of each bubble representing the percentage of chimeras observed in female samples.

### RNA-seq Analyses

RNA samples were extracted from myeloid differentiated cells and cleaned up with RNeasy Kit (Invitrogen). mRNA library preparation and NovaSeq PE 150 were performed by Novogene. The quality of raw data was assessed using FastQC^47^ compiled with MultiQC^48^. Adaptors and low-quality reads were filtered using with a sliding window quality filter^49^. After passing QC, reads were aligned to the human genome (ensembl GRCh38.v110) using Kallisto^50^ with the GC bias flag. K-means clustering was performed using IDEP^32^ with an elbow plot to determine the optimal cluster number.

### Clinical Sample Collection

The use of human clinical blood samples was approved by the IRB committee of the University of Virginia (protocol #13310). Blood samples obtained from the Department of Pathology at the University of Virginia were de-identified. Buffy coats were separated from a total of 1,351 whole blood samples using 2 ml of blood collected in sodium citrate buffer for centrifugation at 4°C and 3000 g for 15 min.

The COVID-19 study was reviewed and approved by the Institutional Review Board of Tongji Hospital, Tongji Medical College, Huazhong University of Science and Technology, China (TJ-IRB20200405). All enrolled patients provided signed informed consent and all blood samples were collected for the rest of the standard diagnostic tests, with no additional burden to the patients. Blood samples from 90 COVID-19 patients without selected comorbidities were collected from Tongji Hospital and Union Hospital of Huazhong University of Science and Technology, Xiangyang Central Hospital, Hubei University of Arts and Sciencxe, and Hubei Dazhong Hospital of Chinese Traditional Medicine between 19 February and 26 April 2020. The patients were classified into four groups according to disease severity as described in Wu P. et al^51^. The ethylenediaminetetraacetic acid disodium salt (EDTA-2Na)-anticoagulated venous blood samples were separated by centrifugation at 3000 rpm for 7 min at room temperature after standard diagnostic tests.

### RNA Extraction

RNA was extracted from buffy coats with TRIzol reagent (Invitrogen) following the manufacturer’s instructions. RNA samples were examined for quantity and quality using Nanodrop (Thermo Scientific), and reverse-transcribed via random primers using the Verso cDNA synthesis kit (Thermo Scientific). All 1,252 clinical blood samples passed QC, with OD ratios of 260nm/280nm, and 260nm/230nm, falling within 1.6-2.0, and levels of an internal control GAPDH within two standard deviations of the mean value.

### RT-PCR and Sanger Sequencing

Primer pairs for PCR were designed using Primer3 (Whitehead Institute for Biomedical Research). A Step One Plus Real-Time PCR System (Applied Biosystems) was used to perform SYBR Green based RT-qPCR assay. Relative RNA levels were calculated using the ΔΔCt method. Gene expression was normalized to that of the housekeeping gene, *GAPDH* (glyceraldehyde 3-phosphate dehydrogenase). Amplified products were separated using agarose gel electrophoresis. Proper size product bands were purified using the PureLink Quick Gel Extraction Kit (Invitrogen) and sent for Sanger sequencing (Genewiz). Primer sequences are listed in Table S1.

### Chromosome Conformation Capture

3C was performed as described in Cope N. et al^52^. Human peripheral blood mononuclear cells (PBMCs) were isolated by Ficoll-Paque (GE Healthcare) density centrifugation from buffy coats (American Red Cross) and stored in liquid nitrogen. After thawing, 10 million PBMCs were fixed in 1.5% formaldehyde for 10 min. Cross-linking was quenched with 0.25 mM of glycine for 10 min at room temperature, followed by two washes with PBS. Cells were resuspended in 10 ml lysis buffer (50 mM Tris–HCl pH 8, 150 mM NaCl, 0.5% NP-40, 1% Triton X-100, protease inhibitor cocktail (Sigma)) for 30 min on ice. Cell lysates were centrifuged at 2,000 rpm for 5 min at 4°C to isolate the nuclei. Nuclei were then resuspended in 500 μl of 1× NEBuffer3 (NEB) containing 0.3% SDS and incubated for 1 h at 37°C, followed by 1 h incubation after adding 10% Triton X-100 (final concentration 1.8%). Nuclei were digested using 800 U of BamHI (for 5’-5’ loop) or BglII (for junction loop) (NEB) on a shaker overnight at 37°C. Digestion was quenched by adding 10% SDS (final concentration 1.6%) and incubated for 25 min at 65°C. The digested nuclei were washed and resuspended by 1 x T4 ligase buffer (NEB). Nuclei were then ligated for 4 h at room temperature by adding 1% Triton X-100, 1 mM ATP, and 4000 U T4 DNA ligase (NEB). The ligation reaction was stopped by adding 0.5 M EDTA (final concentration 10 mM). The ligated chromatin was followed by proteinase K (final concentration 100 ug/ml) digestion at 65°C overnight and then treated with 0.5 ug/ml RNase A for 1 h at 37°C water bath. The DNA was extracted by phenol-chloroform and chloroform and precipitated by 100% ethanol and sodium acetate. DNA pellets were then washed by 70% ethanol and resuspended in 200 μl H2O. 2 μl of 3C DNA was used for qPCR. Unidirectional primers were used to detect ligation interaction frequency. 3C qPCR analysis was performed as described in Naumova N. et al^53^. Primes efficiency was normalized by BAC library and the interaction frequency was normalized against anchor site ligation frequency. Primers for each digestion site were listed in Table S2.

### Chromatin Immunoprecipitation

10 million PBMCs were fixed in 1.5% formaldehyde for 10 min. Cross-linking was quenched with 0.25 mM of glycine for 10 min at room temperature, followed by two washes with PBS. Cells were resuspended in 1 ml lysis buffer (5 mM PIPES pH 8, 85 mM KCl, 0.5% NP-40, protease inhibitor cocktail (Sigma)) for 10 min on ice. Cell lysates were centrifuged at 5,000 rpm for 5 min at 4°C and the nuclei was incubated in the nuclei lysis buffer (50 mM Tris-HCl pH 8, 10 mM EDTA, 1%SDS) for 10 min on ice. Nuclei lysates were sonicated with 30s on and 30s off for 20 min in total. Following sonication, nuclei was cleared by centrifugation and subjected to pre-clearance by incubating with Protein A agarose beads (Cell Signaling). Pre-cleared cell lysates were separately incubated with CTCF or non-specific IgG antibodies (Cell Signaling) overnight, followed by incubation with Protein A agarose beads for 2 h and 4 °C. Protein-bound agarose beads were washed sequentially with low salt wash buffer, high salt wash buffer, LiCl wash buffer, and TE buffer. Beads-bound complexes were subjected to elution by IP elution buffer (1% SDS, 0.1 M NaHCO_3_). Eluted complexes were de-crosslinked by 5 M NaCl overnight at 67°C. All elutes and inputs were degraded in protein/RNA degradation buffer (EDTA, Tris-HCl pH6.5, proteinase K, RNase A). The DNA was extracted by phenol-chloroform and chloroform and precipitated by 100% ethanol and sodium acetate. DNA pellets were then washed by 70% ethanol and resuspended in 60 μl H2O. 2 μl of ChIP DNA was used for qPCR. Percentage of input for each IP or IgG was calculated. Primers for each CTCF binding sites were listed in Table S3.

### CRISPR-Mediated Reorganization of Chromatin Loop Structure

CLOuD9 was performed as described in Morgan S. et al^29^. Briefly, gRNAs targeting each desired anchor site were designed and cloned into *S. aureus* (CSA) and *S. pyogenes* (CSP) constructs. gRNA sequences were listed in Table S4. The plasmids were transfected into HEK293T cells through GeneCellin (Bulldog Bio) following manufacturer’s protocol. 48 h after transfection, cells were selected in growth medium with puromycin (1 μg/ml) and hygromycin (25 μg/ml). Cells were kept in selection media for at least 1 week and were maintained in selection media for the duration of downstream experiments. For dimerization treatment of CLOuD9-transduced cells, 1 mM Abscisic Acid (Fisher Scientific) (or an equivalent volume of DMSO for controls) was added to the culture medium. Medium was changed with fresh abscisic acid or DMSO daily. After a 3-day treatment, cells were collected by trypsin digestion, followed by either 3C analysis or RNA extraction.

### Cell Culture

HEK293T cells were maintained in DMEM medium (Gibco), supplemented with 10% fetal bovine serum (VWR), and 1% Pen/Strep solution (Gibco). Cells were cultured at 37°C in 5% CO2 humidity.

Cryopreserved CD34+ primary human progenitors were purchased from FredHutch and thawed following their instruction. CD34+ cells were maintained in Iscove modified Dulbecco medium (Gibco), supplemented with 20% BITS 9500 (StemCell Technologies), 2 mM l-glutamine (Gibco), 1% Pen/Strep solution (Gibco), and 1-thioglycerol (Sigma). Undifferentiated CD34+ cells were expanded for four days in prestimulation medium by adding 100 ng/ml stem cell factor (SCF; PeproTech), 100 ng/ml FLT3-ligand (PeproTech), 100 ng/ml TPO (PeproTech), and 20 ng/ml IL-3 (PeproTech). For myeloid differentiation, expanded CD34+ cells were cultured in basal medium with 20 ng/μl GM-CSF (Biolegend) and 1 x CC100 (StemCell Technologies) consisting of cytokines SCF, FLt3-ligand, IL-3, and IL-6.

### shRNA Transduction

Oligo DNA sequence of shUC1 and shUC2 were cloned into pLKO.1 plasmid vector. shRNA sequences were listed in Table S4. The plasmid was packaged into lentivirus with packaging constructs pCMV and pMD6.G. Briefly, 750,000 HEK293T cells per well were seeded into a six-well plate. 24 h after seeding, plasmids were transfected into HEK293T cells using GeneCellin (Bulldog Bio) following manufacturer’s protocol. After 24 h, media was changed to viral production media opti-MEM (Gibco). After 24 h, viral production media was collected and spun at 500 g for 5 min to remove cell debris. RetroNectin (Takara) was used to coat six-well plate following manufacturer’s protocol. For transduction, 1 ml viral supernatant was preloaded to the RetroNectin coated plate and incubated at room temperature for 30 min. The viral supernatant was discarded prior to adding cells. 500,000 pre-stimulated CD34+ cells were suspended in 1 ml viral supernatant and 1 ml prestimulation medium with 5 μg/ml protamine sulfate (Sigma). The cells mixture was added to the RetroNectin coated plate and incubated at 37°C in 5% CO2 humidity for 2 h. The plate was then spun at 2,000 rpm for 90 min and incubated at 37°C in 5% CO2 humidity overnight. After 24 h, cells were changed with 1 ml fresh prestimulation medium and 1 ml viral supernatant. After 24 h, cells were selected in prestimulation medium with puromycin (1.5 ug/ml) for 3 days. Successfully transduced cells were selected and initiated with myeloid differentiation.

### Cell Sorting

20 million PBMCs were washed in RPMI media twice before being resuspended in 1 ml of sterile PBS with 5% Fetal Bovine Serum (FBS) in a sterile FACS tube. Antibodies were added to stain surface markers for 30 min at 4°C (CD3 PE, Life Technologies; CD8 FITC, Life Technologies; CD11b AF700, Life Technologies CD19 PerCP-Cy5.5, BioLegend; C33 BV605, BioLegend; CD56 BV711, BD). After 30min, cells were washed twice with PBS+5% FBS then resuspended in buffer with DAPI (1:25,000, Sigma) for FACS sorting. An Influx Cell Sorter (Becton Dickinson) was used to sort cells into subsets, using PBS with 20% FBS as the collection buffer.

### Flow Cytometry

Myeloid differentiated cells were harvested on Day3 and Day5. Cells were washed in PBS and resuspended in 100 μl of PBS+5% FBS. Antibodies were added to stain surface markers for 30 min at 4°C (CD33 BV711, BD; CD11b APC-Cy7, BD; Zombie Violet, BioLegend). After 30min, cells were washed with PBS+5% FBS and fixed in 16% paraformaldehyde. Cells were washed and resuspended in 400 μl of PBS+5% FBS. An Attune flow cytometer (Thermo Fisher) was used to acquire data. Zombie Violet was used to gate out live cells. All live cells were gated out based on their FSC-A and SSC-A profile. Single cells were then gated out with FSC-A vs. FSC-H and used for FACS analysis. Analysis was performed using FCS Express software.

### Statistics

All quantitative data are presented as the mean ± standard deviation. We used unpaired two-tailed t-tests or one-way analyses of variance (ANOVA) for variables analysis. p-values < 0.05 were considered statistically significant.

## Supplementary Figure Legends

**Figure S1.**
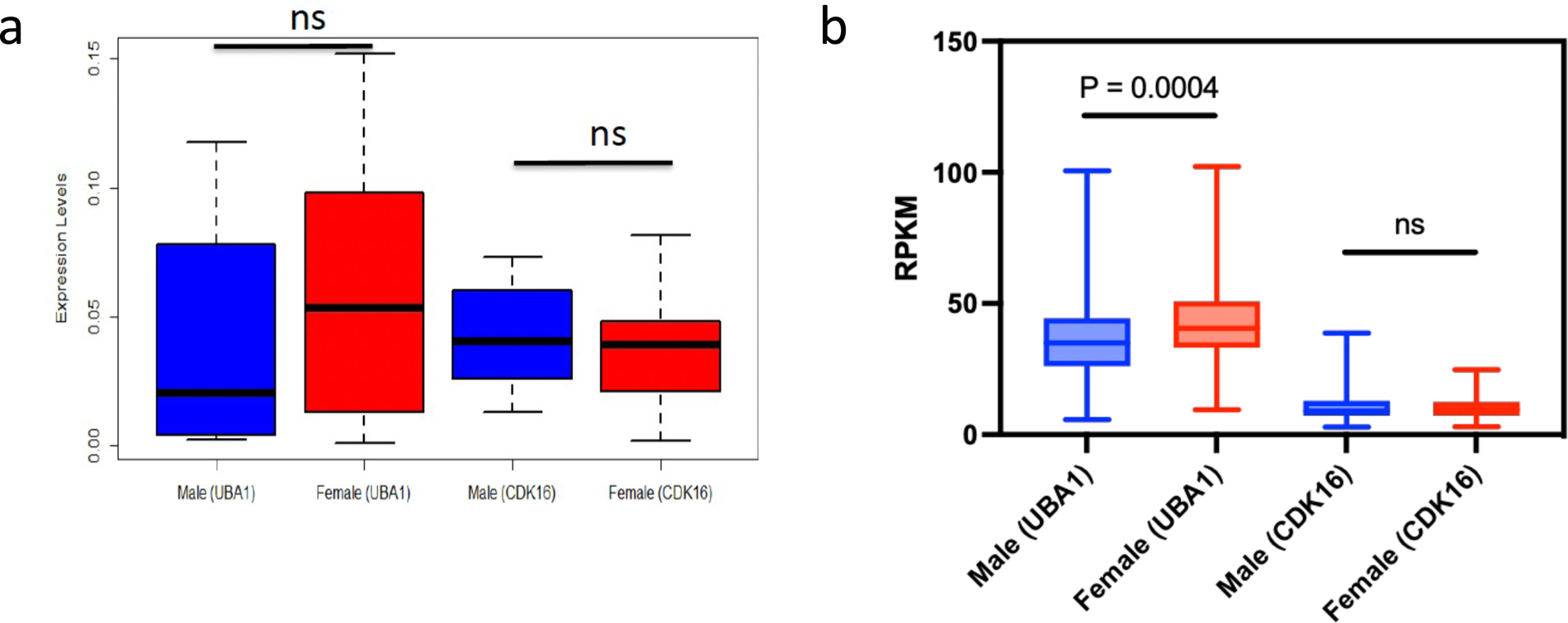
*UBA1* and *CDK16* genes expression in males and females. (a) RT-qPCR detection of parental genes in 15 buffy coat samples. The transcripts were normalized to an internal control *GAPDH.* No statistically significant difference was observed. (b) Comparison of RPKM for *UBA1* and *CDK16* from GTEx whole blood RNA-seq data between 248 males and 144 females.

**Figure S2.**
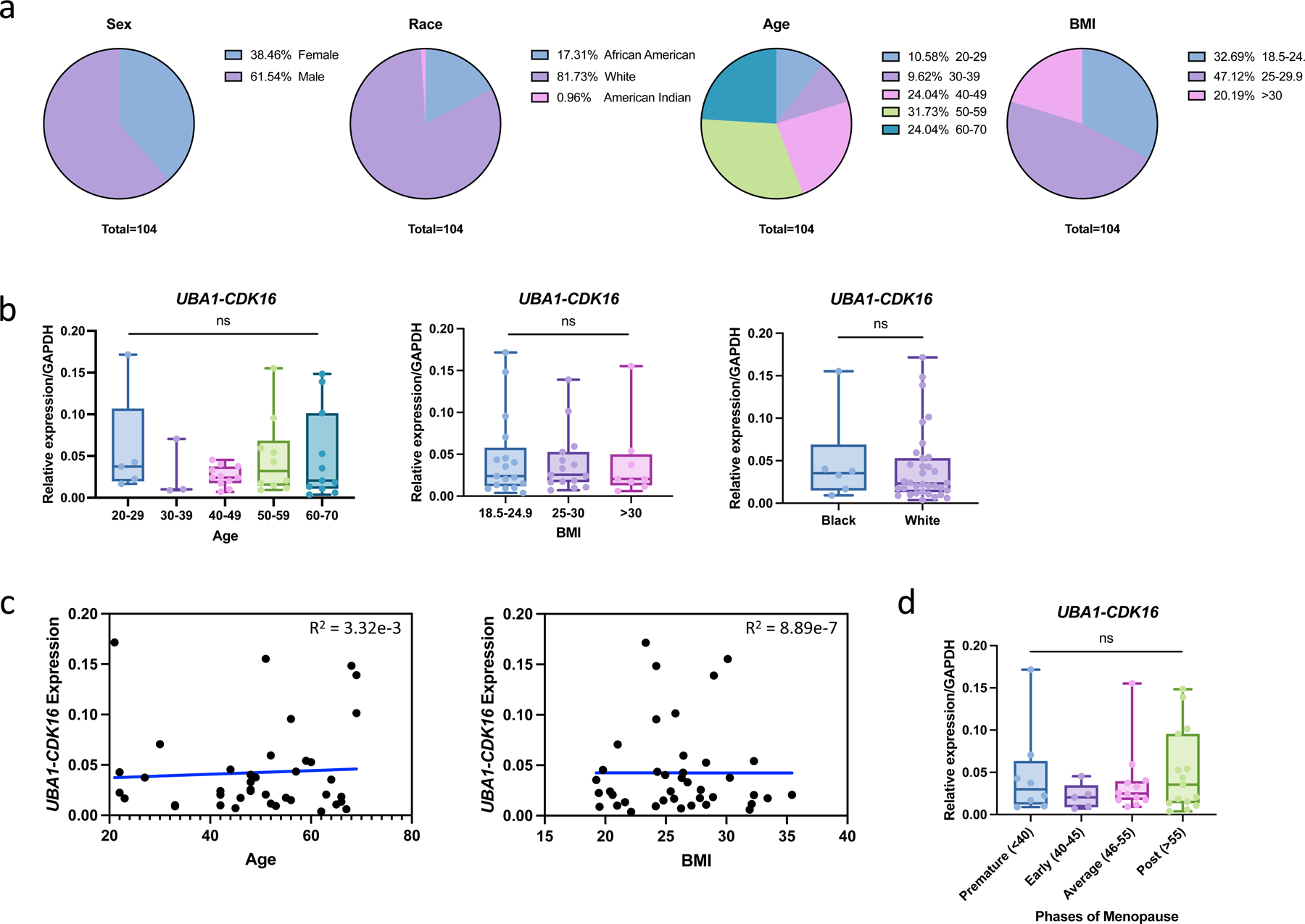
*UBA-CDK16* expression in GTEx blood samples. (a) Distribution of samples in sex, race, age, and BMI. (b) RT-qPCR detection of *UBA1-CDK16* expression in age, BMI, or race groups. (c) Linear regression model of *UBA1-CDK16* expression with age or BMI. R^2^ indicated the non-significant correlation. (d) RT-qPCR detection of *UBA1-CDK16* expression in different phases of menopause. The transcripts were normalized to an internal control *GAPDH.* No statistically significant difference was observed.

**Figure S3.**
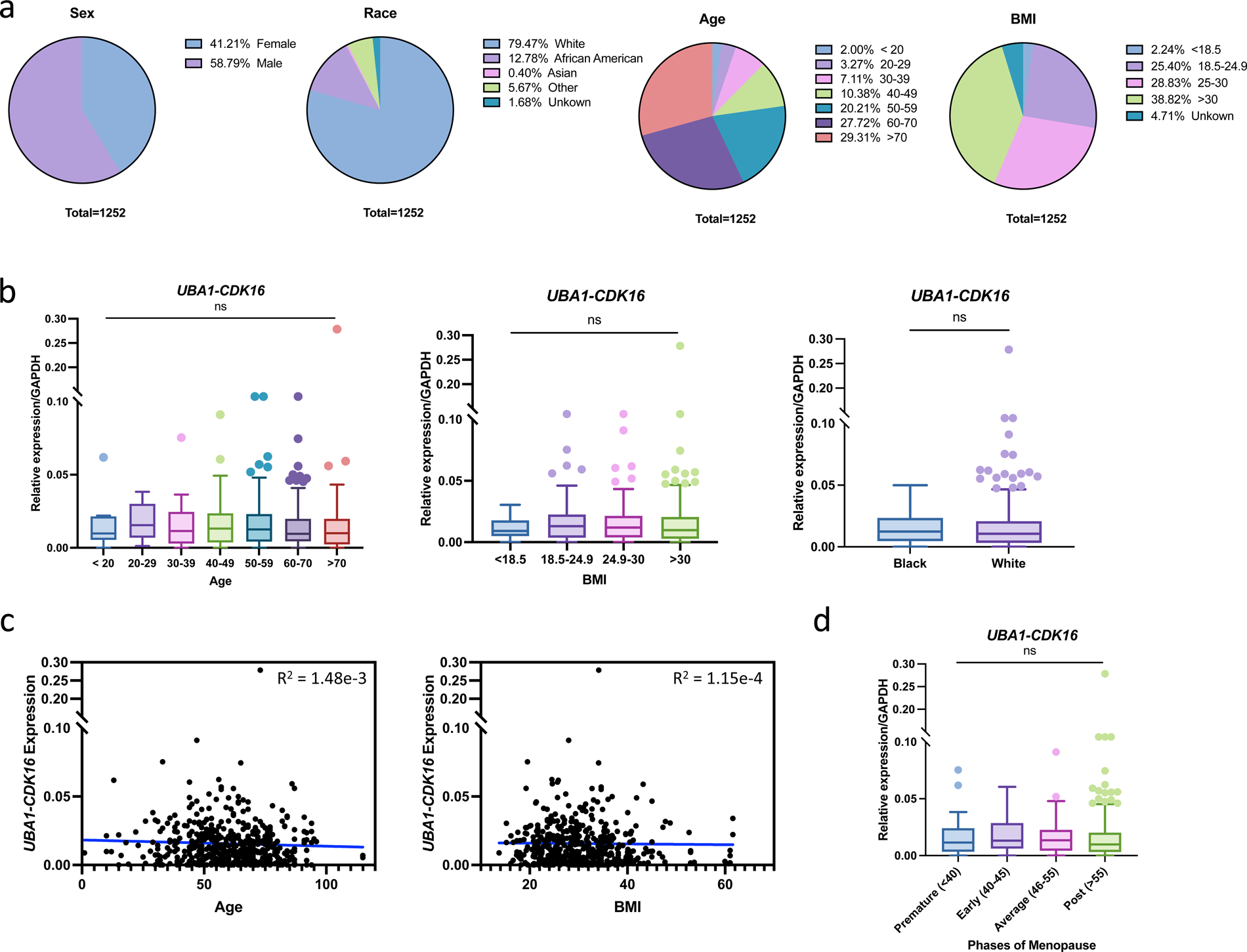
*UBA-CDK16* expression in UVa clinical blood samples. (a) Distribution of clinical samples in sex, race, age, and BMI. (c) RT-qPCR detection of *UBA1-CDK16* expression in age, BMI, or race groups. (d) Linear regression model of *UBA1-CDK16* expression with age or BMI. R^2^ indicated the non-significant correlation. (e) RT-qPCR detection of *UBA1-CDK16* expression in different phases of menopause. The transcripts were normalized to an internal control *GAPDH.* No statistically significant difference was observed.

**Figure S4.**
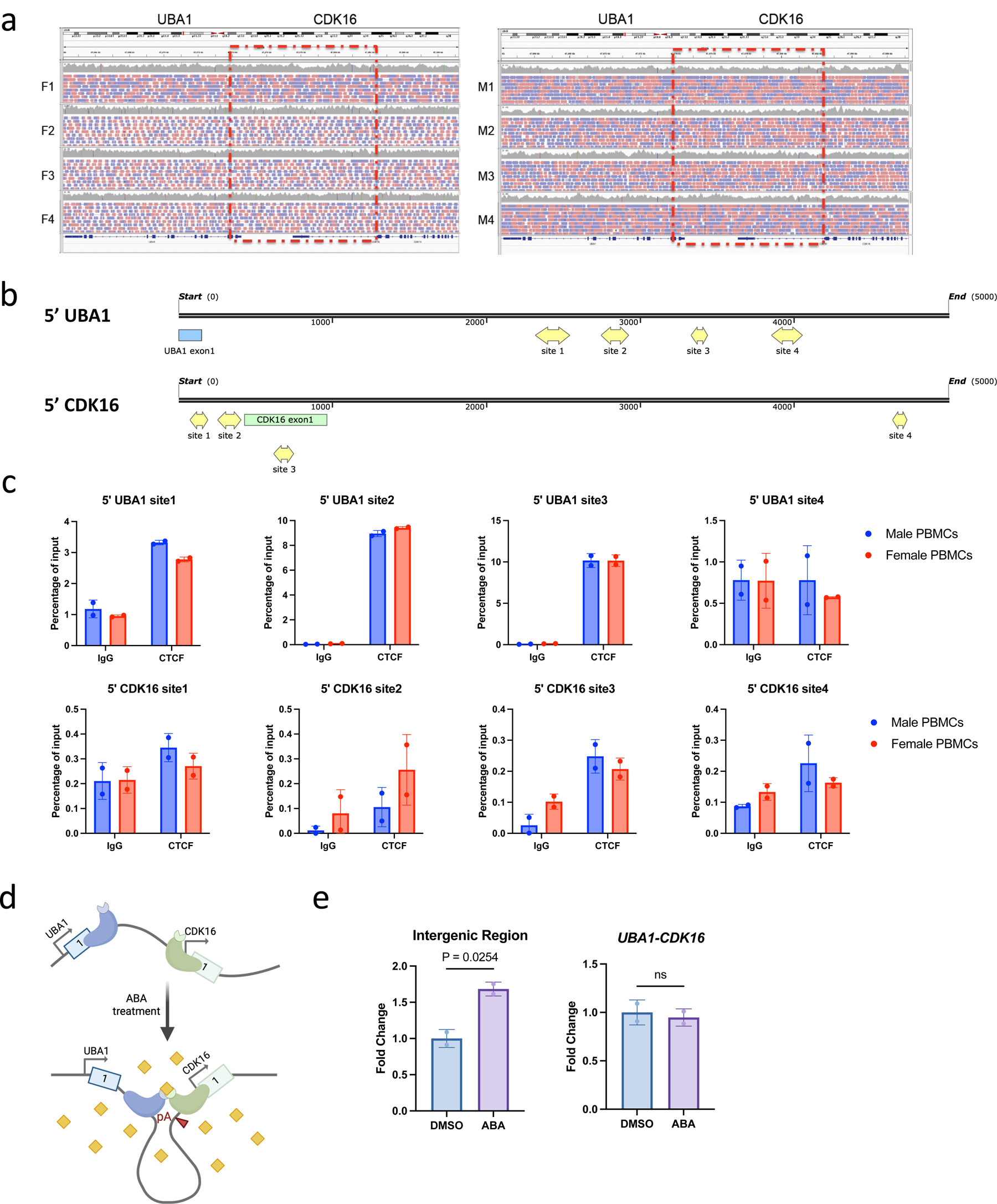
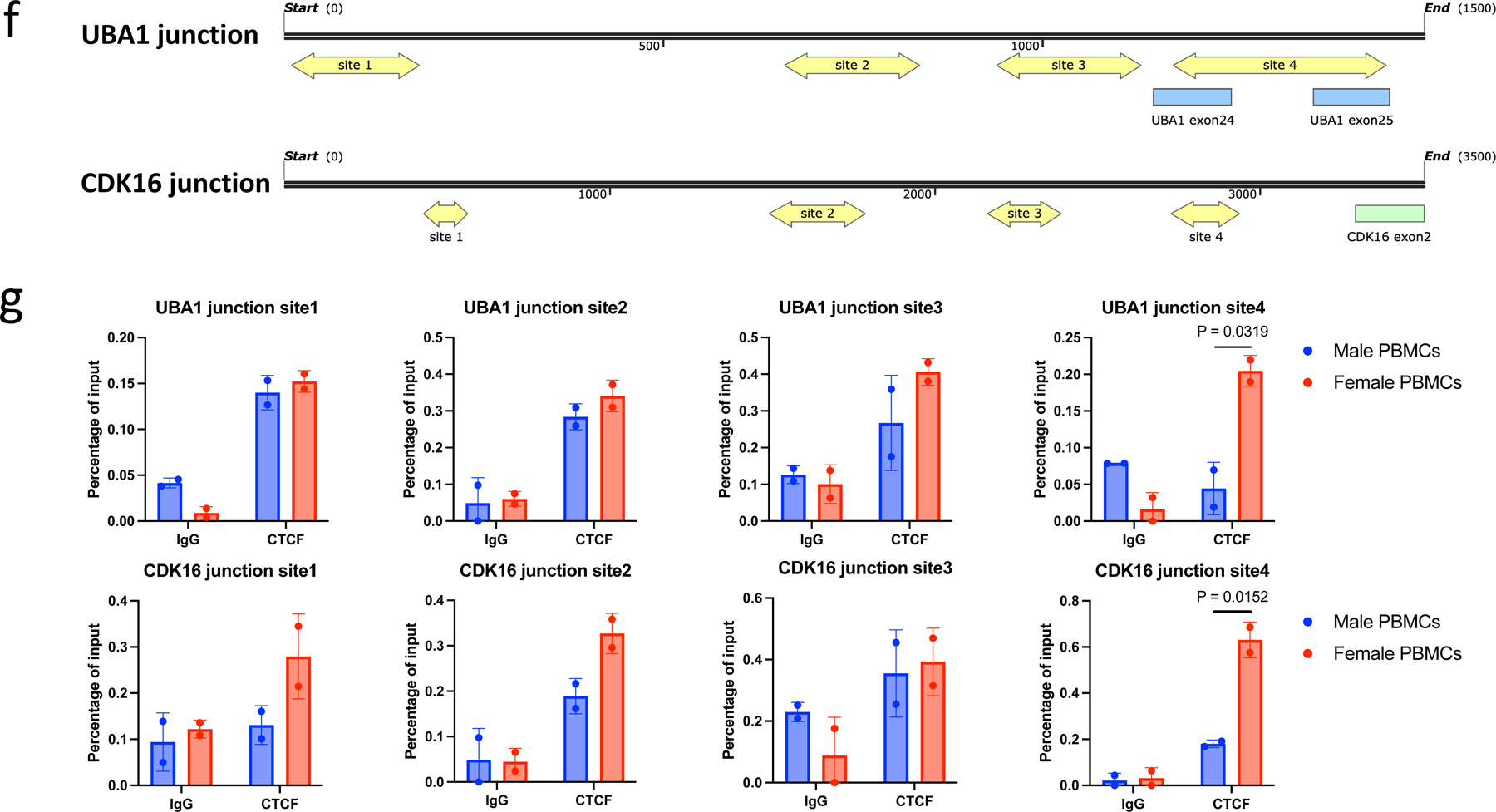
*UBA1-CDK16* is a product of cis-SAGe through alternative splicing. (a) Whole genome sequencing of GTEx whole blood samples showed no evidence of interstitial deletion between *UBA1* and *CDK16* in four females and four males. (b) Schematic of CTCF binding sites at 5’ regions of *UBA1* and *CDK16*. CTCF binding sites were predicted by the CTCFBSDB 2.0 database. (c) CTCF ChIP-qPCR in male and female PBMCs at each binding site. IgG antibody was used as control. (d) CLOuD9 experimental design with sgRNAs targeting the 5’ regions of *UBA1* and *CDK16*. 5’-5’ chromatin loop was induced by abscisic acid (ABA) treatment. (e) cis-SAGe-qPCR revealed a significant increase of precursor readthrough mRNA after ABA treatment. The relative expression level was normalized against *CDK16* using primers targeting exon 1, and fold change was normalized against samples treated with DMSO. No statistically significant change was detected on mature chimeric RNA *UBA1-CDK16* revealed by RT-qPCR. (f) Schematic of CTCF binding sites at junction regions of *UBA1* and *CDK16*. (g) CTCF ChIP-qPCR in male and female PBMCs at each binding site. IgG antibody was used as control.

**Figure S5.**
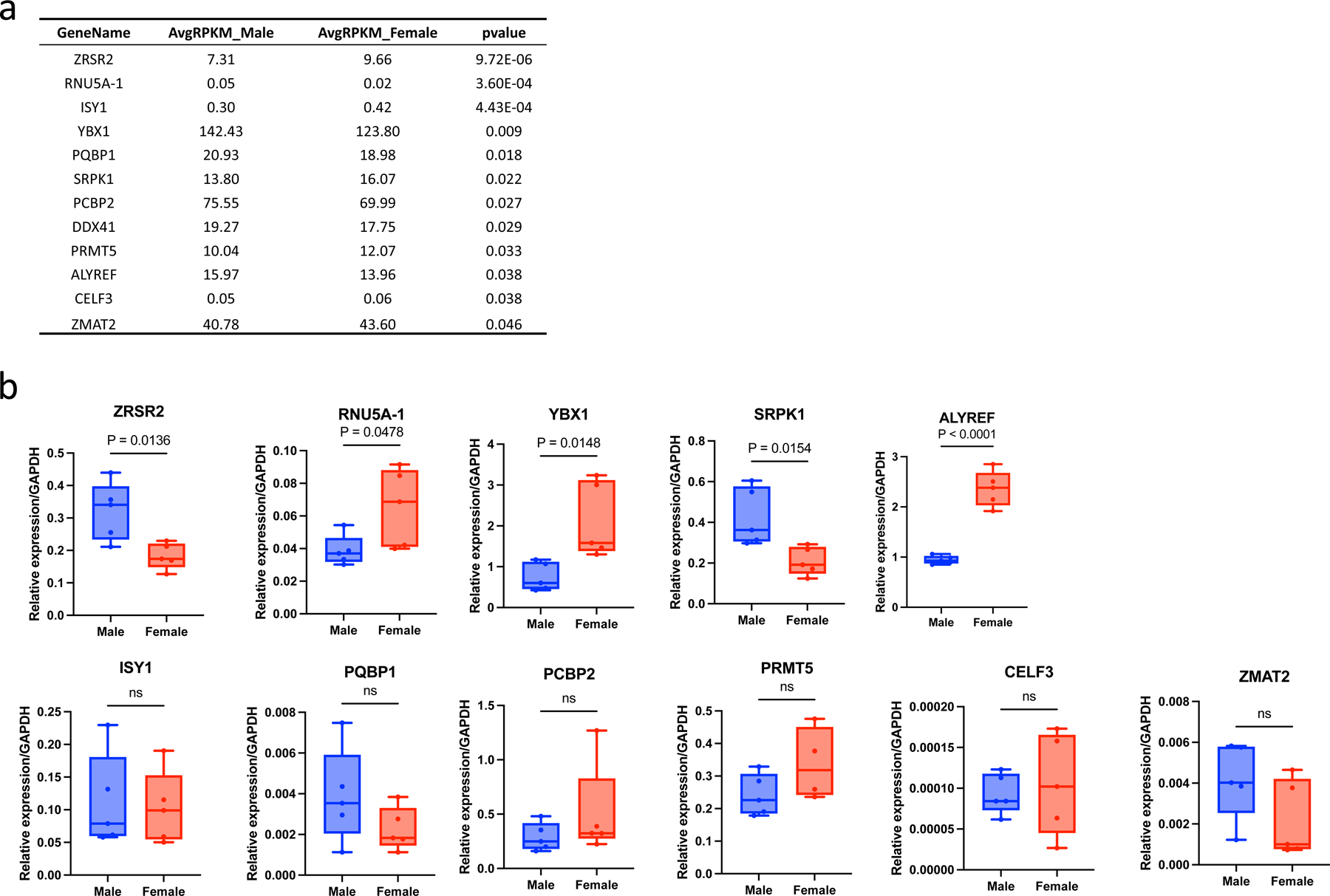
Expression of sex-differential splicing factors in clinical blood samples. (a) List of splicing factors with significant sex difference, compared between average RPKM from GTEx male and female whole blood RNA-seq. Statistical difference was determined with a p-value cut off < 0.05. (b) RT-qPCR validation of splicing factors in five male and five female buffy coats. The transcripts were normalized to an internal control *GAPDH*.

**Figure S6.**
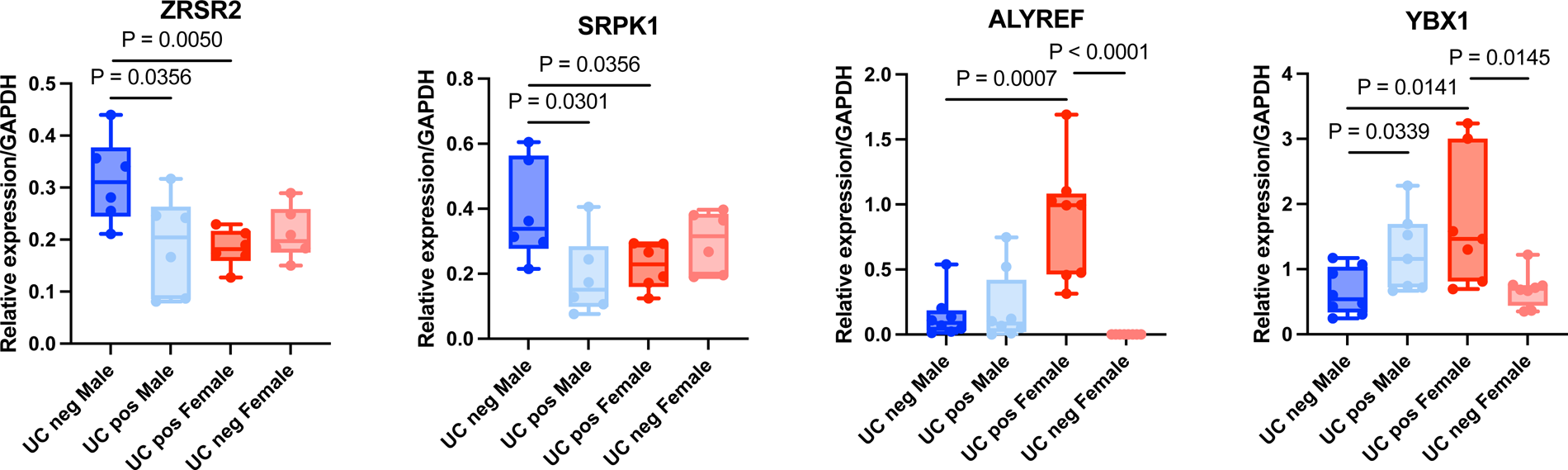
Expression of sex-differential splicing factors in outlier clinical blood samples. RT-qPCR detection of various splicing factors in outlier clinical blood samples from Fig. 1h. The transcripts were normalized to an internal control *GAPDH*.

**Figure S7.**
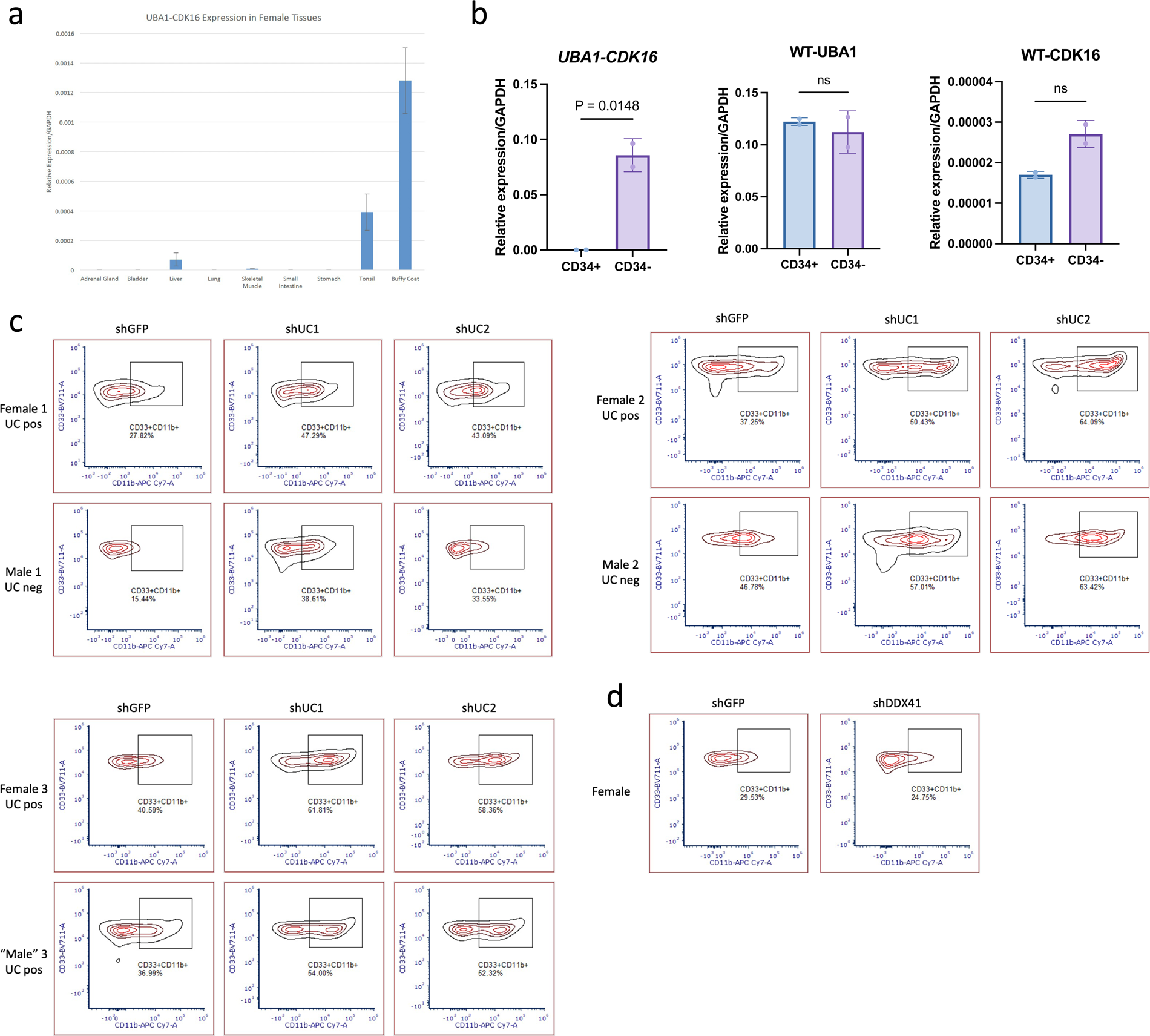
*UBA1-CDK16* inhibits CD11b expression during myeloid differentiation. (a) RT-qPCR detection of *UBA1-CDK16* expression in human female tissues, including adrenal gland, bladder, liver, lung, skeletal muscle, small intestine, stomach, tonsil, and buffy coat from whole blood. Levels of the chimeric transcript were normalized to that of *GAPDH*. (b) Chimeric RNA *UBA1-CDK16* and parental genes expression in CD34+ and CD34-cells. CD34+ cells were isolated from PBMCs using CD34 microbeads, while CD34-cells comprised all remaining cells. A statistically significant difference was observed in chimeric RNA expression between CD34+ and CD34-cells. (c,d) Flow cytometry analysis of single cells gated for CD33+CD11b+ myeloblasts in three pairs of CD34+ cells donors.

**Figure S8.**
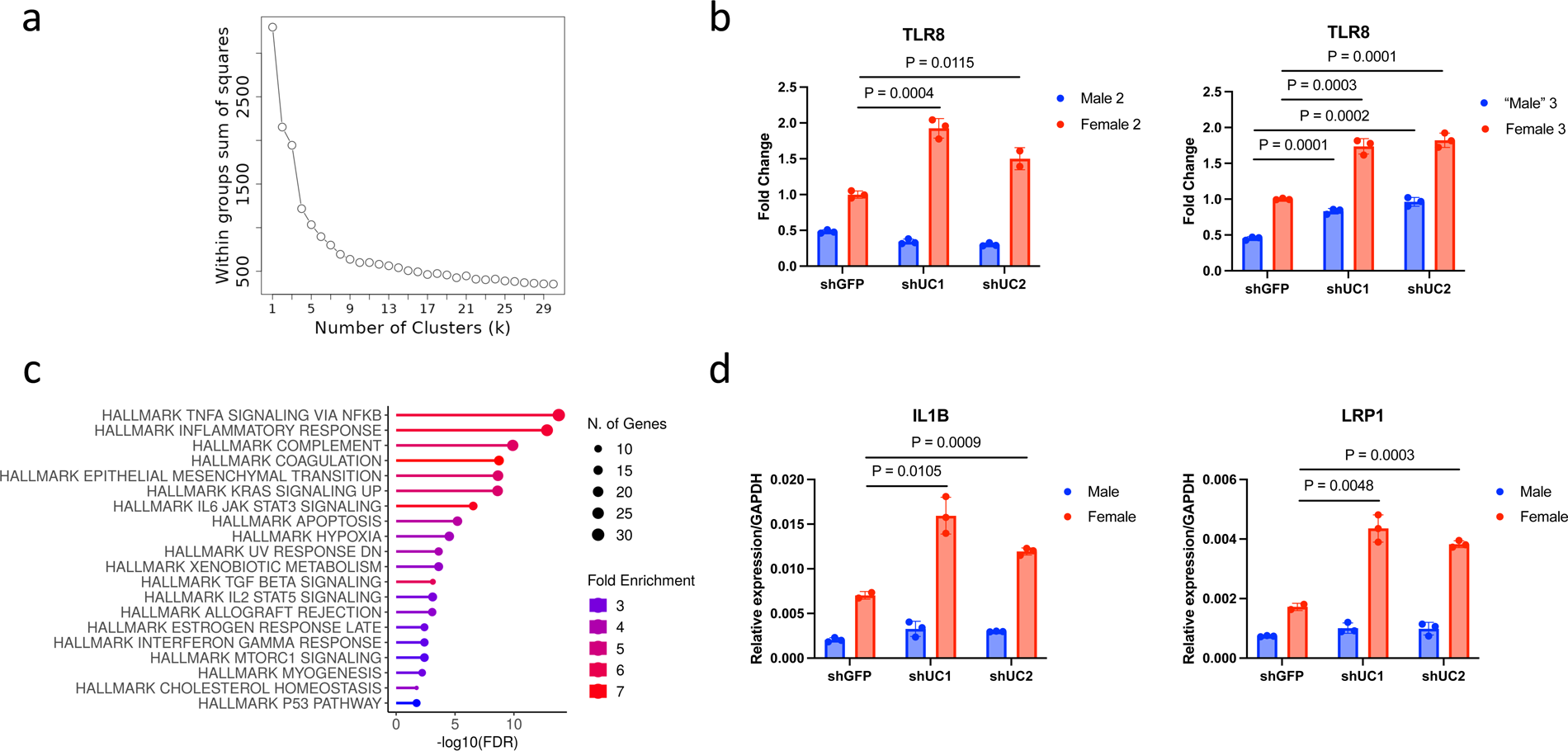
Discovery of *UBA1-CDK16* downstream targets through RNA-seq analysis. (a) The elbow method conducted by iDEP estimated the optimal cluster number used for k-means clustering. (a) RT-qPCR detection of *TLR8* expression in RNA samples extracted from day 3 of myeloid differentiated cells, obtained from the second and third pairs of CD34 cells donors. (c) Enriched hallmark pathways of differentially expressed genes in cluster A. The enriched pathways were ranked by FDR. (d) RT-qPCR validation of *IL1B* and *LRP1* in RNA samples extracted from day 3 of myeloid differentiated cells.

**Figure S9.**
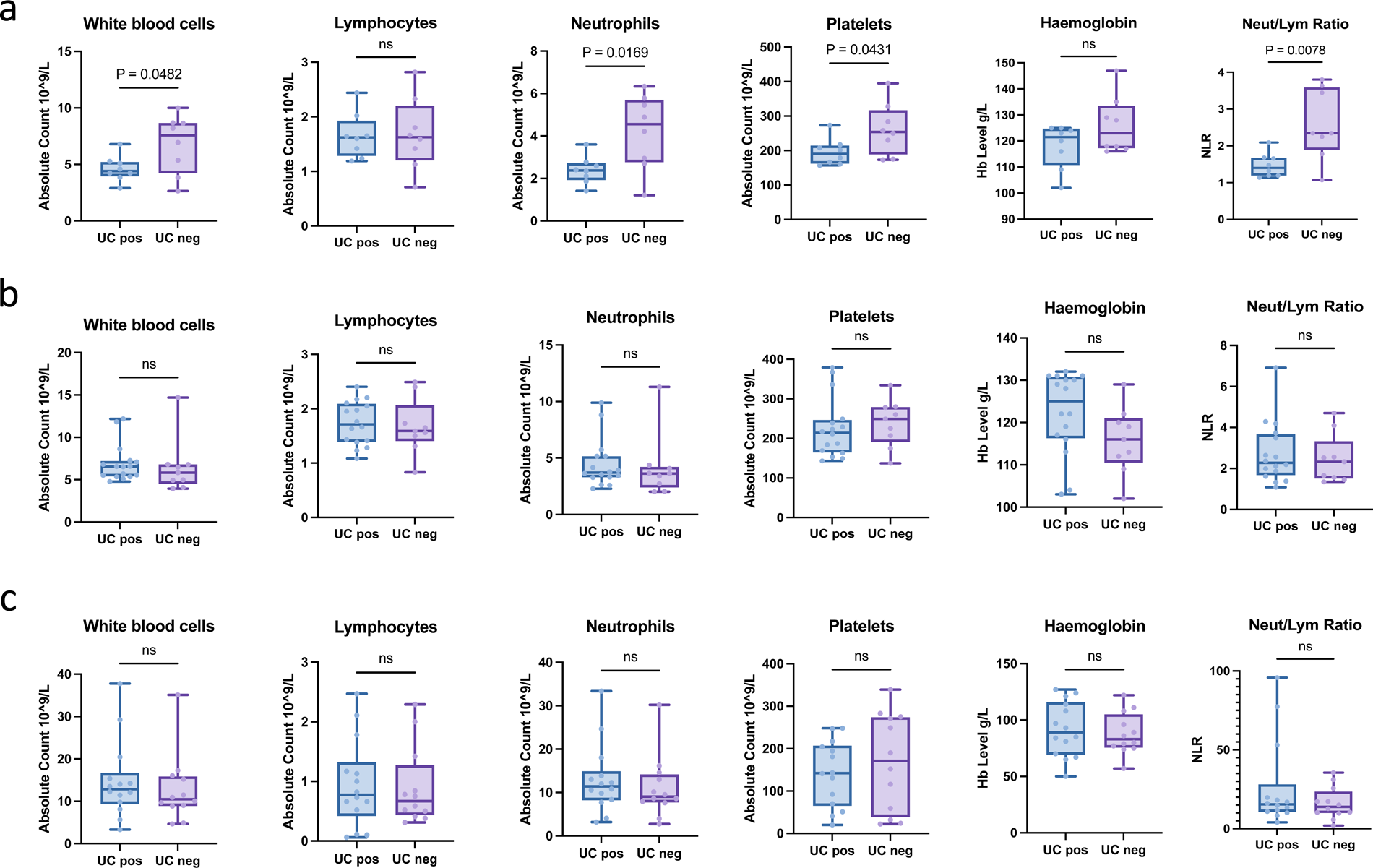
Blood cell counts in COVID-19 female patients. White blood cells, Lymphocytes, Neutrophils, Platelets counts, Hemoglobin level, and NLR were detected in (a) mild, (b) severe, and (c) critical COVID-19 female patients. Comparison was made between patients detected positive or negative for *UBA1-CDK16*.

**Figure S10.**
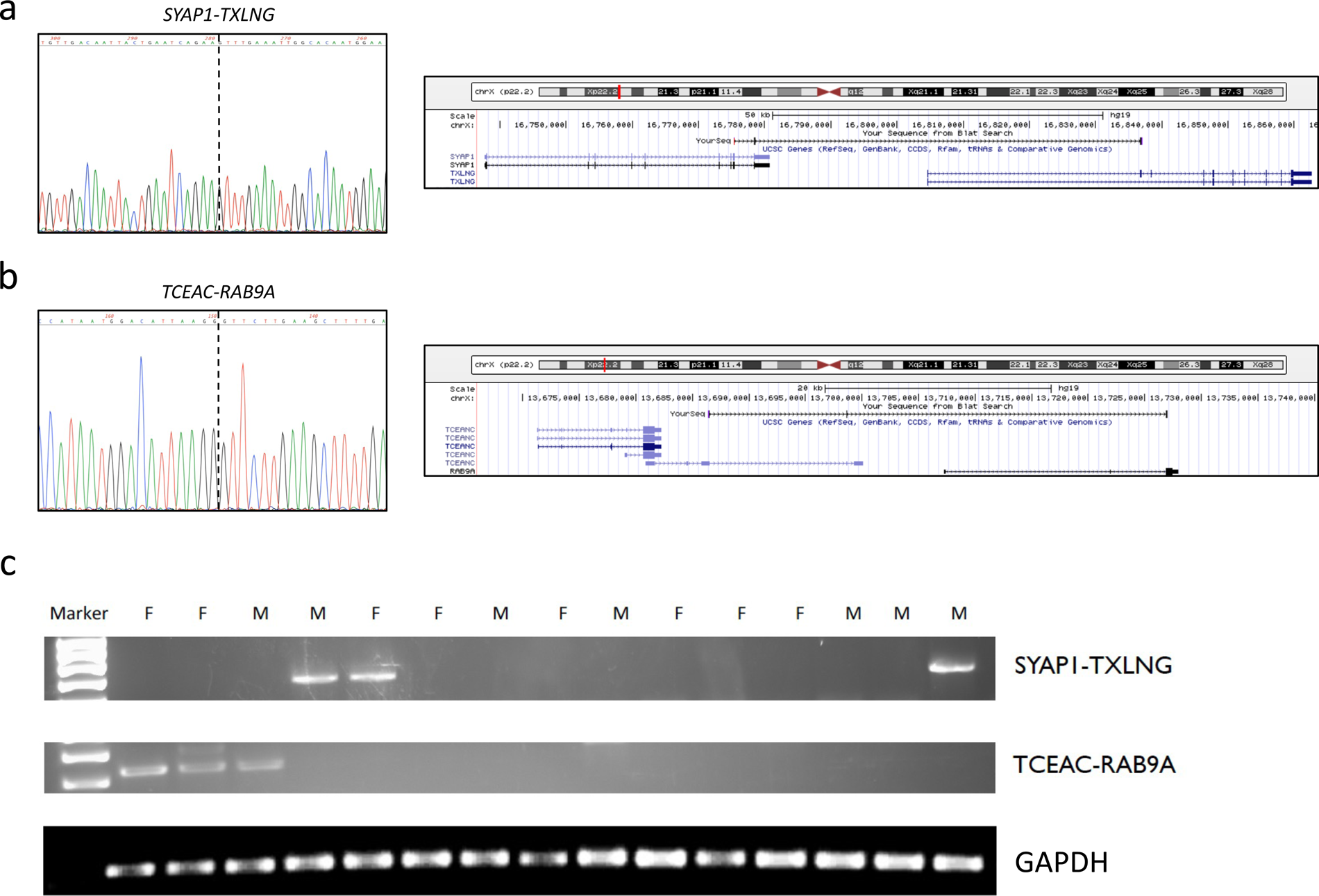
Other chimeric RNAs expressed from escaping zone of X inactivation are not female-biased. X-linked cis-SAGe chimeric RNA, *SYAP1-TXLNG* and *TCEANC-RAB9A*, were amplified by RT-PCR using combinations of primers that anneal to each parental gene. Sanger sequencing of the RT-PCR products is shown on the left, and alignment of the sequence to the hg38 human genome in the UCSC browser is shown on the right. (a) *SYAP1-TXLNG* chimeric RNA involves the third-to-last exon of *SYAP1* joining to the second exon of *TXLNG*. (b) *TCEANC-RAB9A* chimeric RNA involves the second-to-last exon of *TCEANC* and the second exon of *RAB9A*. (c) RT-PCR validation of the expression of the chimeric RNAs in eight female (F) and seven male (M) samples (same as used in Fig. 1c). Gel electrophoresis revealed the presence of the chimeric RNA in both female and male samples, even though many samples showed no expression.

